# Regional specificity of neuronal cell types, functional connectivity, and cell type-specific correlation to brain disorders in the human ventral tegmental area

**DOI:** 10.64898/2026.06.08.730903

**Authors:** Anna S. van Regteren Altena, Bernardo de A.P.C. Maciel, Rachel M. Brouwer, Chen J.R. Jiang, Sara L. Seoane, Ravian L. van Ineveld, Jonne M. Eggink, Tiziana M. Hey, Anne C. Rios, R. Jeroen Pasterkamp, Vanessa Donega, Martijn P. van den Heuvel, Onur Basak, Elly M. Hol

**Author notes:** Correspondence &.

## Abstract

The ventral tegmental area (VTA) in the midbrain is a key reward system hub associated with multiple brain disorders. While rodent studies reveal a diverse neuronal population and distinct connectivity patterns, the architecture of the human VTA remains poorly understood. Here, we generated a transcriptomic reference atlas of the adult human VTA with subregional resolution. We identify multiple neuronal cell types defined by differential transcription factor expression, as well as regionally enriched GABAergic cell types. Functional connectivity analysis with fMRI suggests that this spatial organization corresponds to distinct connectivity profiles. Integration with other human midbrain single-nucleus RNA-seq datasets reveals a VTA-specific combinatorial GABA-dopaminergic cell type and regional differences among dopaminergic neurons. Finally, we find multiple associations between neuronal cell types and genetic risk for psychiatric disorders and body mass index. Altogether, this study provides a foundation for mechanistic and therapeutic studies into VTA-related diseases.

## Main

The ventral tegmental area (VTA) is a key midbrain region in the reward system containing dopaminergic (DA) neurons that project to different brain areas^1^. Single-cell RNA sequencing (scRNA-seq) studies in rodents showed the complexity of the neurotransmitter-based neuronal cell types in this area^2–6^. In addition to DA neurons, the VTA contains GABAergic, glutamatergic (GLUT), and combinatorial neurons that secrete multiple neurotransmitters^1,2,4,7–13^. Within the VTA, the inputs from and outputs to other brain areas are heavily regulated by this heterogeneous neuronal population^7,13–15^. The different neuronal cell types are spatially organized within the mouse and rat VTA and have region-specific connectivity, which is involved in different behaviors^9,16–22^. For instance, there are large differences in neuronal cell type composition and connectivity patterns across the medial-lateral axis of the mouse VTA, with DA-GLUT neurons in the medial VTA projecting to the prefrontal cortex (PFC) and DA neurons in the lateral VTA to subregions of the nucleus accumbens (NAc)^9^, and GABA neurons in the dorsal VTA of the rat that mainly project to the anterior cingulate cortex^19^. We hypothesize that the human VTA is regionally organized as well, but we lack experimental evidence on the identity, distribution, and connectivity of its cells.

Multiple studies of the human midbrain have emerged in the past couple of years, revealing its cellular composition at the single-cell and spatial level. However, a single-cell dataset specific to the VTA is still lacking^23–28^. This is in part because most of these studies on the midbrain focused on the substantia nigra (SN), or the midbrain was taken as a whole, making it difficult to distinguish the VTA cells. Also, the human VTA has a complex 3D structure and poorly defined boundaries^7^. Unlike the adjacent SN and red nucleus (RN), the human VTA is practically embedded in white matter, which makes the anatomical identification of its subnuclei difficult^7^. Furthermore, within the midbrain the main focus has been on profiling the DA population in the SN, because of its differential sensitivity to degeneration in Parkinson’s disease (PD)^28^. The differences between VTA and SN DA neurons in the human brain are largely unknown, and unraveling these could help in understanding cell type- and brain region-specific implications in brain diseases.

Here, we generated a reference atlas with single nucleus RNA sequencing (snRNA-seq) of subregions of the adult human VTA to determine neuronal subclasses and assessed their spatial distribution. In addition, we used functional magnetic resonance imaging (fMRI) to investigate region-specific functional connectivity within the human VTA that could be linked to the regional specificity of cell types. Next, we integrated our VTA snRNA-seq with other adult human midbrain atlases to uncover a VTA-specific neuronal cell type and the diversity of DA cell types within the VTA and SN. Finally, using this integrated atlas, we discovered disease-associated neuronal cell types in the midbrain through associations with genome-wide association study (GWAS) summary statistics, with non-DA neurons being associated with multiple psychiatric disorders. Overall, this study characterizes regional heterogeneity of VTA-specific neuronal cell types and connectivity in the human brain, which contributes to the understanding of the human reward system and its related diseases.

## Results

### Transcriptome-defined cellular diversity of the adult human Ventral Tegmental Area

To investigate the cell type diversity of the human VTA across its subregions, we performed snRNA-seq on VTA tissue from six non-diseased donors (**Fig. 1a**, **Suppl. Fig 1**, **Methods**, **Suppl. Table 1**). First, the borders of the VTA with adjacent regions were identified. We divided the VTA into three subregions to study the regional cell type organization, based on myelin presence and immunofluorescent staining for tyrosine hydroxylase (TH) that shows the spatial organization of DA neuron soma. We defined a lateral, medial, and centrally located dorsomedial (which we refer to as central) segment that connects the medial and lateral subregions, in accordance with the differential cell type composition and connectivity along the medial-lateral axis of the mouse VTA^18^, and the proposed localization of subnuclei in the adult human VTA^7^ (**Suppl. Fig. 1**, **Methods**). An unbiased approach was taken: all DAPI-positive nuclei were sorted after isolation, including neurons, glia, and non-neural cells, and we retrieved a total of 85,254 nuclei after quality control (**Fig. 1b**, **Suppl. Fig. 2**, **Methods**). Oligodendrocytes formed the most abundant cell class, followed by astrocytes and microglia (**Fig. 1b**). Classification using the Walktrap algorithm identified 68 clusters, of which neurons displayed the highest number of subclusters (**Fig. 1c, 1d**, **Methods**). Following a hierarchical cell type annotation proposed by the Allen Institute for Brain Science (**Methods**), cell classes were determined using known cell marker genes; i.e., *MAP2* for neurons, *AQP4* for astrocytes, *CX3CR1* for microglia, *CNP* for oligodendrocytes*, CSPG4* for oligodendrocyte precursor cells (OPCs), *PDGFRB* for perivascular cells including pericytes and smooth muscle cells, *PTPRB* for endothelial cells, *CD163* for perivascular macrophages (PVM), and *SKAP1* for T-cells (**Fig. 1d**). Our snRNA-seq data revealed the composition of neuronal and glial cell types in the adult human VTA and highlighted a heterogeneous neuronal population together with a large glial network.

**Fig. 1.**
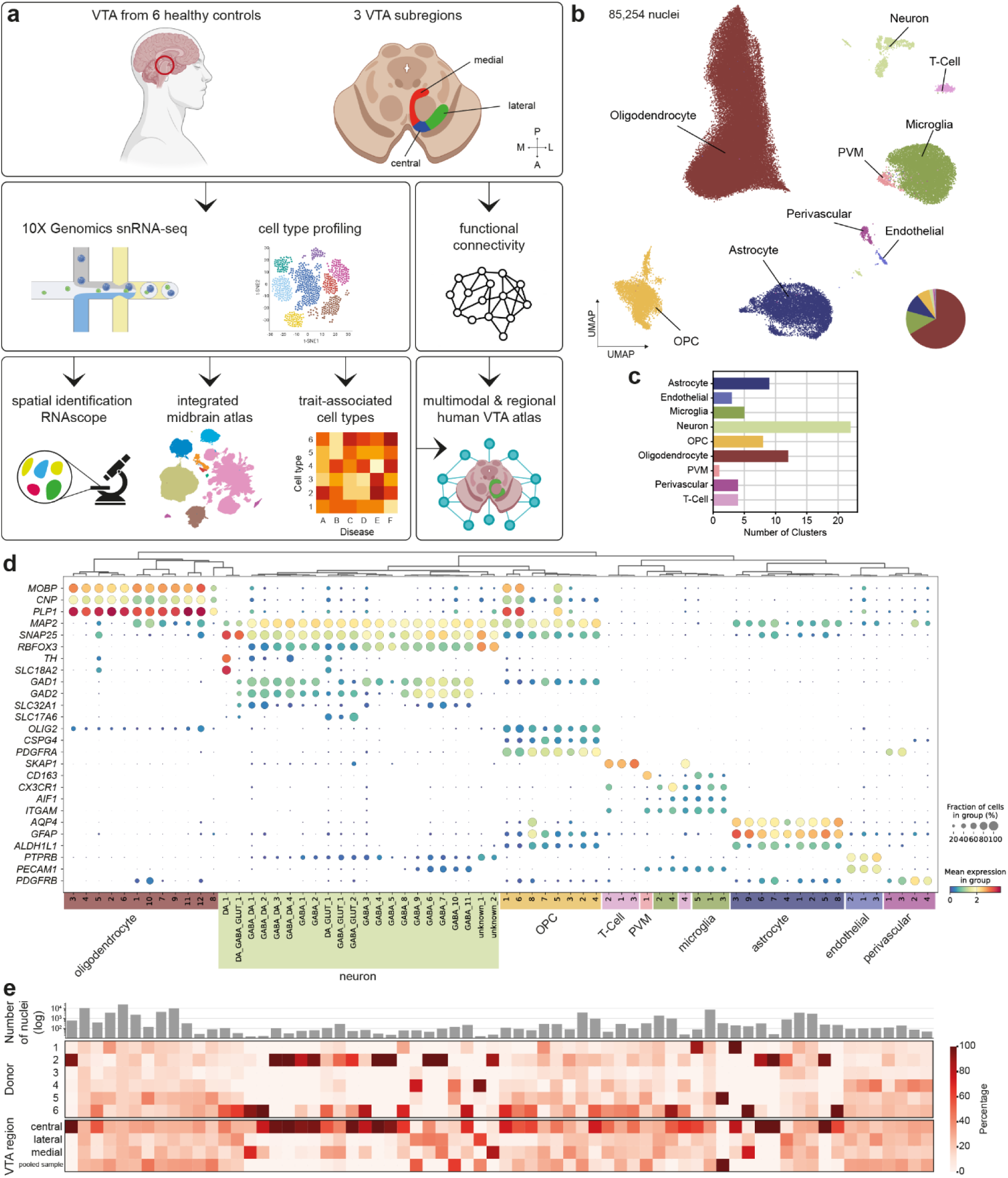
snRNA-seq as a molecular basis for cell type mapping in the human VTA. **a**, Schematic overview of the workflow of this study. 12 samples were collected from six non-demented donors (four males and two females, mean age 80) and three VTA regions (red, medial; blue, central; green, lateral), on which snRNA-seq was performed to identify cell types and further downstream analysis (**Suppl. Table 1**, **Suppl. Table 2**). **b**, UMAP of the 85,254 isolated nuclei and identified cell classes in the human VTA with snRNA-seq. **c**, Number of clusters per cell class. **d**, Dot plot of cell class-specific gene expression in the 68 hierarchically clustered cell types, indicated with the mean normalized expression in the fraction of cells expressing a particular gene. **e**, Nuclei number per cluster in log scale, and contribution to the cell classes by the different donors and VTA regions.

### Neuronal diversity in the adult human VTA is highly complex

To characterize the extent of neuronal diversity in the adult human VTA, we further analyzed the neuronal cell class consisting of 1,632 nuclei. Clustering using the Walktrap algorithm revealed 22 neuronal clusters, which were analyzed separately (**Fig. 1d, Suppl. Fig. 3, Methods**). We grouped them into neurotransmitter-based neuronal subclasses, based on their hierarchical relationships and the RNA expression of proteins necessary for specific neurotransmitter synthesis and release machineries (**Fig. 2a**). These include *TH* (TH; tyrosine hydroxylase) and *SLC18A2* (VMAT2; vesicular monoamine transporter 2), necessary for dopamine synthesis and release, *SLC17A6* (VGLUT2; vesicular glutamate transporter 2) for the release of glutamate, and *GAD1* (GAD67; glutamate decarboxylase 67)*, GAD2* (GAD65; glutamate decarboxylase 65), and *SLC32A1* (VGAT; vesicular GABA transporter) for the synthesis and release of GABA (**Fig. 2b, Suppl. Fig 4**). This resulted in the identification of seven neuronal subclasses: DA-only, GABA-only, combinatorial subclasses DA/GABA, DA/GLUT, GABA/GLUT, DA/GABA/GLUT, and an unknown *FAT2*-expressing neuronal population that lacked expression of neurotransmitter-specific genes but did express pan-neuronal markers *MAP2* and *RBFOX3* (NeuN) (**Fig. 2a-b**, **Suppl. Fig. 4a-b**). All subclasses were present in each VTA subregion, with the DA/GABA subclass enriched in the central VTA (**Fig. 2c**). We confirmed this regional aspect with *in situ* hybridization of *TH*, *GAD1*, and *SLC17A6* on a tissue section of three non-diseased donors using RNAscope^29^ (**Fig 2d**, **Suppl. Fig. 6**, **Suppl. Fig. 7**). Additionally, we identified a GLUT-only neuronal subclass with *in situ* hybridization that we could not annotate in our snRNA-seq data, underscoring the importance of complementary methods to identify cell types in human post-mortem brain tissue. We also observed subclass-specific spatial patterns in every subclass, for example, the triple combinatorial DA/GABA/GLUT nuclei located closely around the red nucleus.

**Fig. 2.**
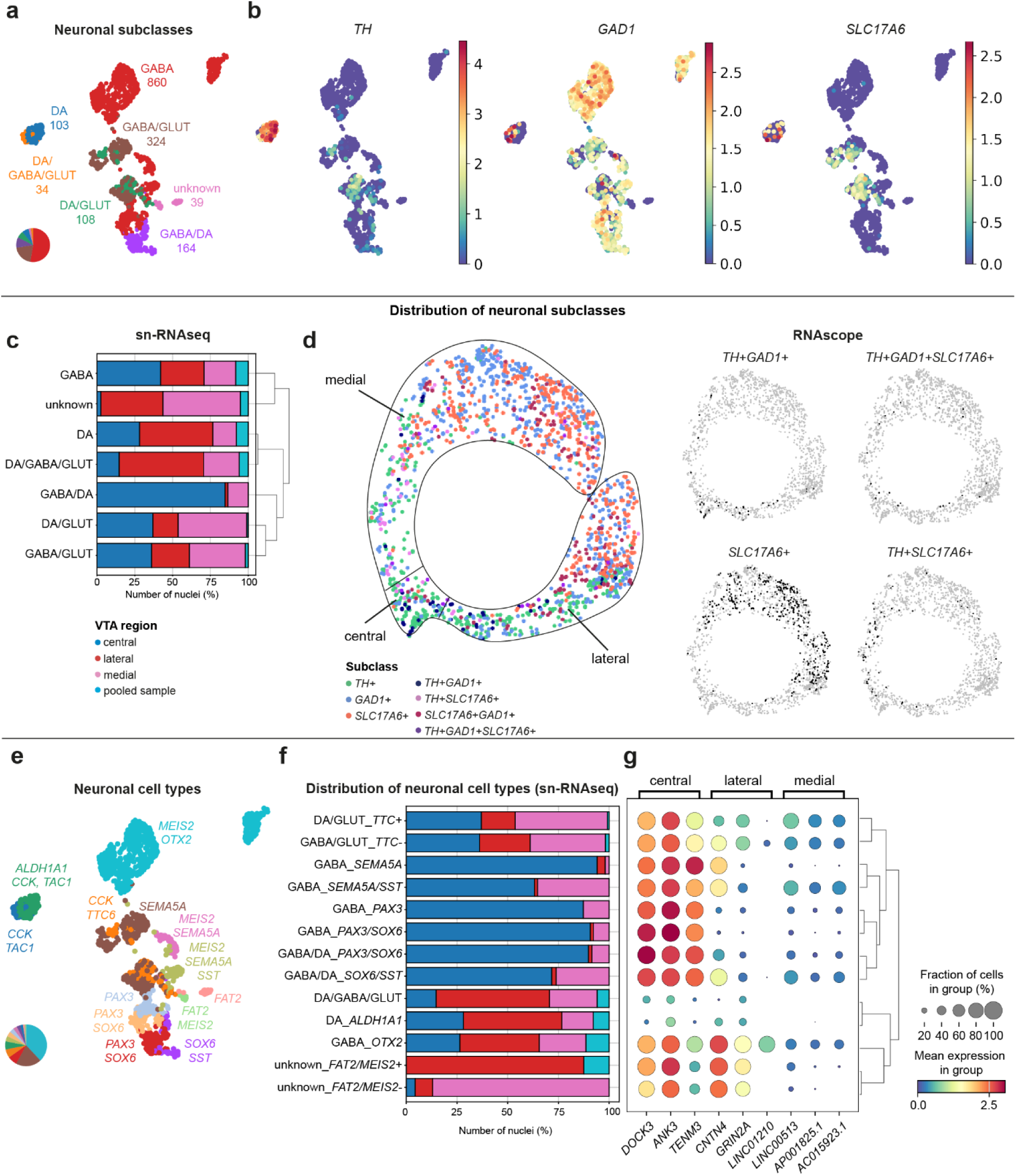
Neuronal subclass and cell type diversity in the adult human VTA. UMAP of seven neuronal subclasses identified on neurotransmitter-related gene expression in the snRNA-seq atlas. **b**, UMAPs of normalized gene expression of neurotransmitter-related genes *TH*, *GAD1,* and *SLC17A6*. **c**, Distribution of the neuronal subclasses in the VTA subregions using the snRNA-seq data. **d**, Representative spatial localization of the neuronal subclasses in a tissue section of a non-diseased donor using RNAscope, with all subclasses combined (left) and four individual subclasses shown (right). **e**, UMAP of annotated neuronal cell types using marker gene profiles. **f**, Distribution of the neuronal cell types in the VTA subregions. **g**, VTA subregion-specific gene expression in each neuronal cell type, identified with differential gene expression (DGE) analysis.

The neuronal cell class contained one DA-only cluster, which, together with the triple combinatorial DA/GABA/GLUT cluster, did not show *RBFOX3* gene expression, in contrast to all other neuronal clusters (**Suppl. Fig. 4a-b**). RNA expression of neuropeptides *CCK*, *SST,* and *TAC1* was present in different subclasses and their subclusters, indicating co-expression of neurotransmitters and neuropeptides as previously summarized^30^, which adds to the modulatory capacity of the neuronal circuitry in the adult human VTA. Within the GABA neuronal subclass, the snRNA-seq data revealed 11 clusters, which were subdivided into *OTX2+* and *OTX2-* groups based on differential gene expression (DGE) analysis of each cluster (**Suppl. Fig. 4b, 4f**). Within the *OTX2-*clusters, DGE analysis showed marker gene expression of transcription factor and axon guidance genes that further split the groups into annotated cell types, including *MEIS2*, *SOX6*, and *PAX3,* and *SEMA5A*, respectively (**Fig. 2e**, **Suppl. Fig 4**). *OTX2-* GABA neurons originated mostly from the central VTA, whereas *OTX2+* GABA neurons were present in all regions (**Fig. 2f**). All four DA/GABA clusters expressed *SOX6* and were divided into *PAX3*+ and *PAX3*- groups, indicating a transcription factor subdivision within this neuronal subclass. Moreover, most DA/GABA neuronal cell types originated from the central VTA, indicating region-specific neuronal cell types (**Fig. 2c**, **2f**). Next, we investigated the interregional molecular differences. DGE analysis between VTA subregions of all neuronal nuclei revealed region-specific gene expression, such as *DOCK3*, *ANK3* and *TEMM3*, that were highly enriched in the central VTA (**Fig. 2g**). Our analysis also revealed that most cell types expressed these DGEs – except for the DA-only and DA/GABA/GLUT cell types – indicating that region-specific gene expression in the VTA is driven by region-specific cell types. Altogether, molecular-based spatially aware cell type identification using snRNA-seq revealed a heterogeneous neuronal population in the adult human VTA identifying several novel neuronal subclasses and cell types and highlighted regional neuronal differences.

### VTA subregions show specific functional connectivity

The spatial location of VTA neurons correlates with the connectivity specificity with different brain regions, as shown in rodent studies^21^. To study the connectivity profiles of the human VTA, we assessed whether its subregions showed different connectivity profiles (**Methods**). Seed-based connectivity analysis of 181 subjects of the Human Connectome Project^31^ using resting-state fMRI showed that the VTA is functionally connected with the cerebral and cerebellar cortices, as well as with several subcortical structures (**Fig. 3a, 3b, Suppl. Fig. 8**). These included brain regions known to be structurally connected to the VTA in rodent models, such as the prefrontal cortex and nucleus accumbens^32^.

**Fig. 3.**
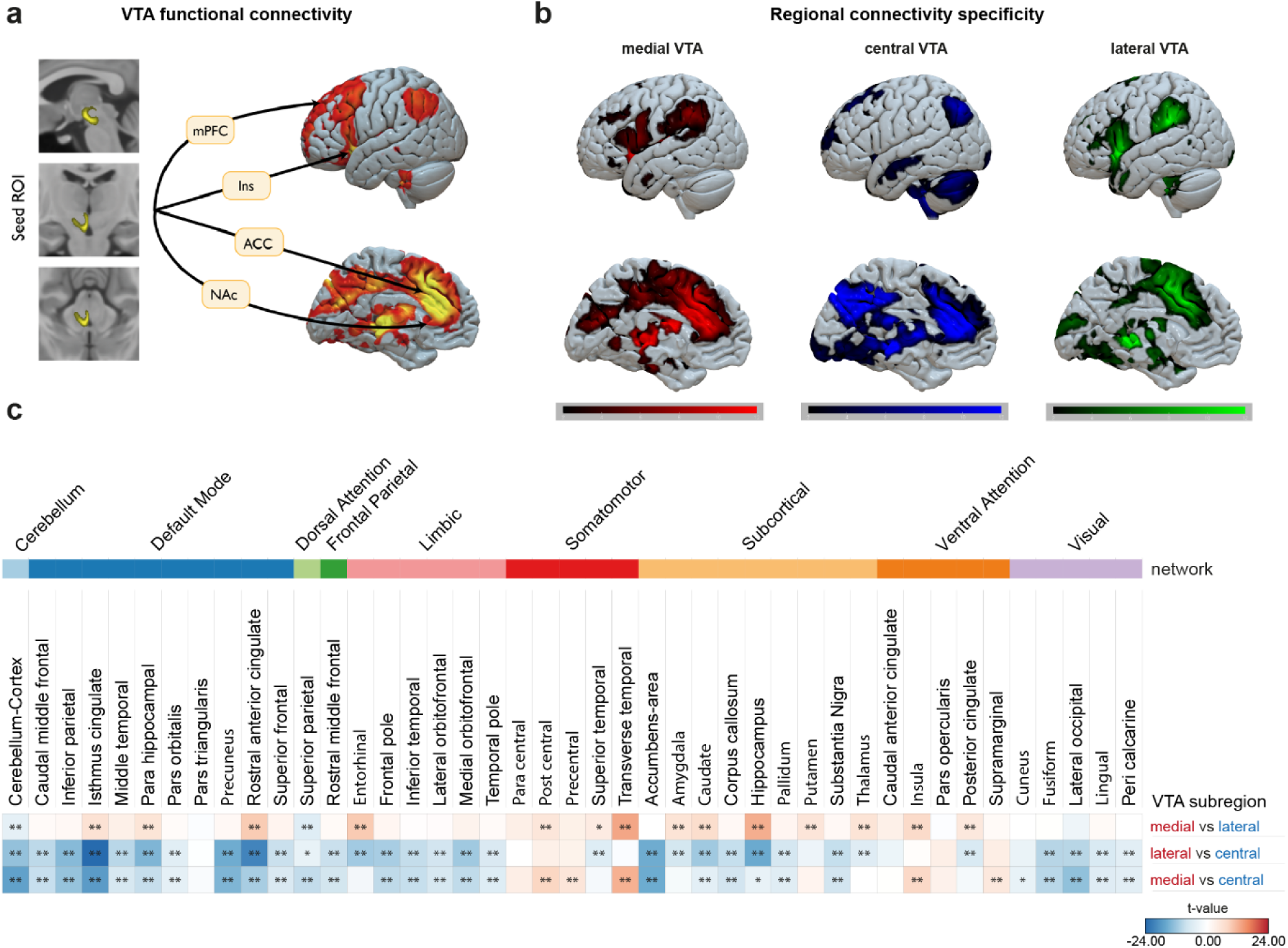
VTA subregions show different connectivity profiles. **a**, Functional connectivity map of the whole VTA. **b**, Group average functional connectivity maps for whole-brain connectivity with the medial, central, and lateral VTA. Color legends show connectivity strength intensities. **c**, Pairwise analyses for region-specific connectivity. Each row shows a different pair of VTA subregion comparisons. The connectivity strength for each comparison is color-coded and shown in t-values, and its significance is indicated by *= p<0.05 and **= p<0.01.

To investigate the specificity in connectivity between VTA subregions, we tested whether central, medial, or lateral subdivisions were more strongly connected with 44 brain regions of interest. Pairwise testing (i.e., medial vs central, medial vs lateral, central vs lateral) revealed that 32 brain areas made significantly stronger connections with the central subdivision of the VTA than with the lateral VTA, and 28 areas made stronger connections with the central than the medial VTA (**Fig. 3c, Suppl. Fig. 9**). Furthermore, there were 20 brain regions for which the medial VTA area had significantly higher connectivity than the central VTA and 13 higher than the lateral VTA. All regions in the somatomotor network displayed significantly higher connectivity with the medial VTA, including the primary auditory, sensory, and motor cortices. The amygdala, a subcortical structure of importance in the reward system^33^, was specifically associated with the medial VTA. Connectivity of the central VTA was significantly stronger with 12 brain regions when compared against the lateral VTA and stronger with 8 when compared to the medial VTA. These included subcortical structures of the reward system such as the nucleus accumbens^34,35^, and the brainstem and cerebellum. The lateral VTA displayed significantly higher connectivity to 4 regions compared to the medial VTA, with the superior parietal lobe showing the strongest association. These results were largely consistent after repeating the analysis using the von Economo MRI-based cortical parcellation (**Suppl. Note 1**, **Suppl. Fig. 10**). These results show that the VTA has both homogeneous and regionally defined connectivity profiles.

### Integrated midbrain atlas reveals VTA-specific neurons in the human midbrain

In the mouse brain, single-cell studies have shown DA neuron diversity in the VTA and its marker genes, revealing multiple DA subpopulations^8,36^. Moreover, multiple DA cell types have been identified in the human SN using snRNA-seq, supporting the possibility of identifying multiple DA cell types in the human brain^28^. Also, the midbrain contains multiple DA regions, including the VTA and SN, which are distinctly susceptible to neurodegeneration^37^. However, the molecular profile of human (DA) neurons in these regions, which could hold information about their susceptibility, is not known yet for the VTA. To determine whether subpopulations of DA cell types and midbrain region-specific gene profiles exist in the human VTA, the number of DA nuclei was increased by integrating our snRNA-seq dataset with three published SN and two midbrain snRNA-seq datasets of the adult human midbrain (**Fig. 4a-b, Suppl. Fig. 11, Methods**)^23–27^. This yielded 157,244 predicted neurons that were originally grouped in 75 clusters (**Suppl. Fig. 12**, **Suppl. Fig. 13**). Further clustering of two *TH*-high clusters resulted in a final total of 89 clusters, which were used for downstream analyses (**Fig. 4c**, **Suppl. Fig. 12**, **Suppl. Fig. 13**, **Methods**). We identified eight neuronal subclasses based on neurotransmitter profiles, which originated from ten dissected midbrain regions (**Fig. 4d-e**, **Suppl. Fig. 10-11**). All DA subclasses, including combinatorial, originated mostly from the SN, and the DA/GABA subclass contained the majority of nuclei that originated from the VTA (**Fig. 4f**). VTA neuronal nuclei clustered together with different midbrain regions, except one DA/GABA cluster (DA/GABA_1) that consisted of 49,59% of VTA nuclei, and 19,4% from the SN-RN region from the Siletti dataset that could be nuclei originating from the VTA (**Fig. 4d-e**, **Fig. 5c-d**, **Fig. 6b**)^24^. This cluster also contained most nuclei from the four DA/GABA clusters from the VTA (**Suppl. Fig. 12c**), indicating that the DA/GABA clusters we identified in the VTA are VTA-specific. To find VTA-specific neuronal gene expression, we performed DGE analysis of neuronal nuclei between midbrain regions. This identified the gene *FBXL7* with highest expression in the VTA, and predominantly in the DA/GABA cluster (**Fig. 4g**). Conversely, highly expressed genes in the SN were broadly expressed across all midbrain regions but largely reflected their high expression in DA-only neurons. In general, most of the region-enriched genes showed expression overlap across multiple midbrain regions, indicating an absence of uniformly region-restricted transcriptional signatures. Partition-based graph abstraction (PAGA) analysis^38^ showed that the VTA is connected with multiple neighboring midbrain regions, indicating continuous rather than sharply segregated transcription profiles across the midbrain (**Fig. 4h**).

**Fig. 4.**
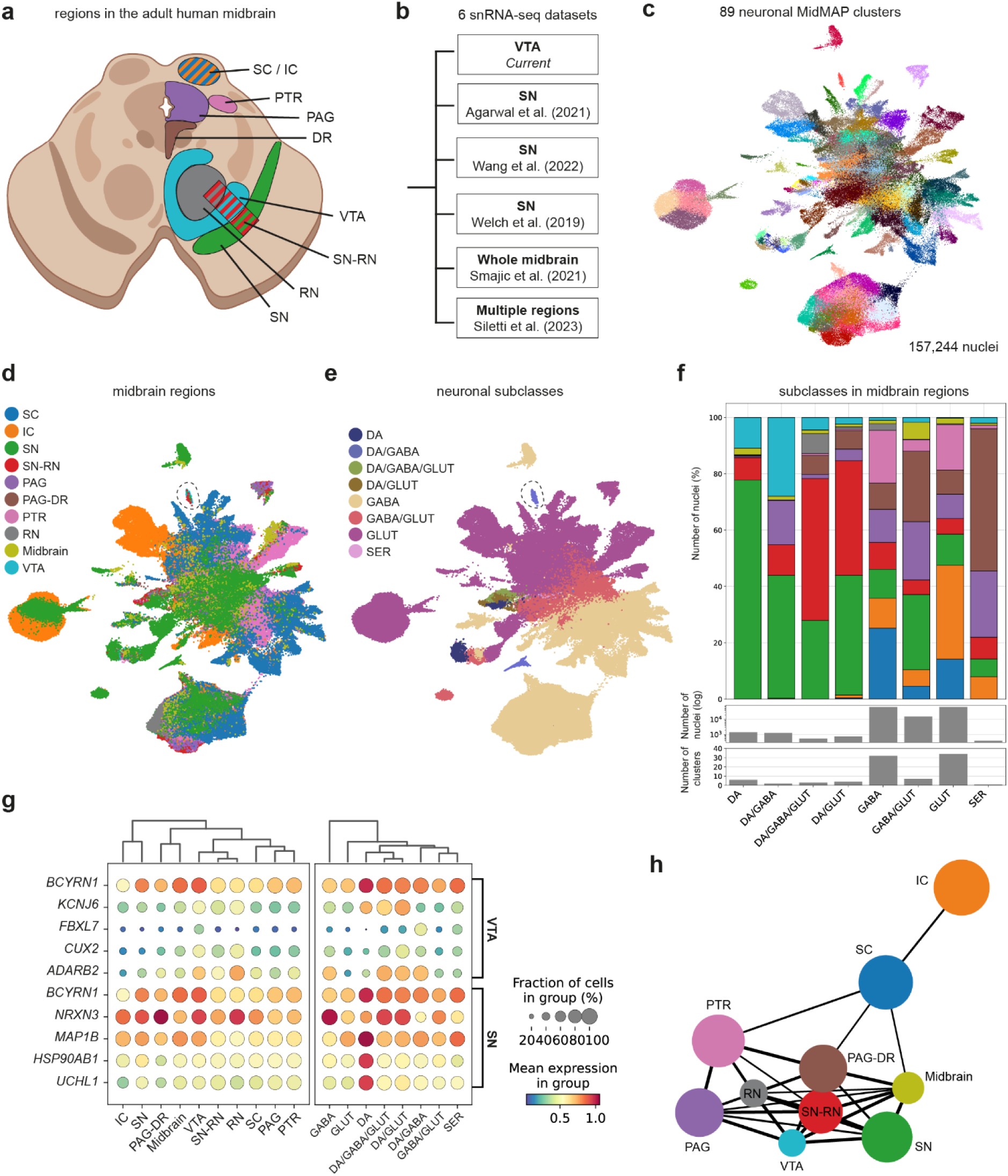
Integrative analysis of adult human midbrain neurons shows a VTA-specific neuronal cell type. **a**, Schematic image of the isolated midbrain regions in a transversal section of the human midbrain. SC, superior colliculus; IC, inferior colliculus; SN, substantia nigra; RN, red nucleus; PAG, periaqueductal grey; DR, dorsal raphe nucleus; PTR, pretectal region; VTA, ventral tegmental area. **b**, Overview of the six atlases that were used for the integration. **c**, UMAP of the 89 identified neuronal clusters of the integrated adult human midbrain atlas, which includes the further clustering of the *TH*-high neuron clusters. **d**, UMAP representation of the different midbrain regions. **e**, UMAP of the neurotransmitter-based identified neuronal subclasses. The cluster circled in **d** and **e** represents the VTA-specific DA/GABA cluster. **f**, Percentage of neurons from each midbrain region, the number of nuclei, and clusters present in each neuronal subclass. The color scheme is the same as in **d**. **g**, Dot plot of differentially expressed genes by the VTA and SN in the different midbrain regions and neuronal subclasses. **h**, Partition-based graph abstraction (PAGA) plot showing the computational connectivity of the clusters between midbrain regions.

**Fig. 5.**
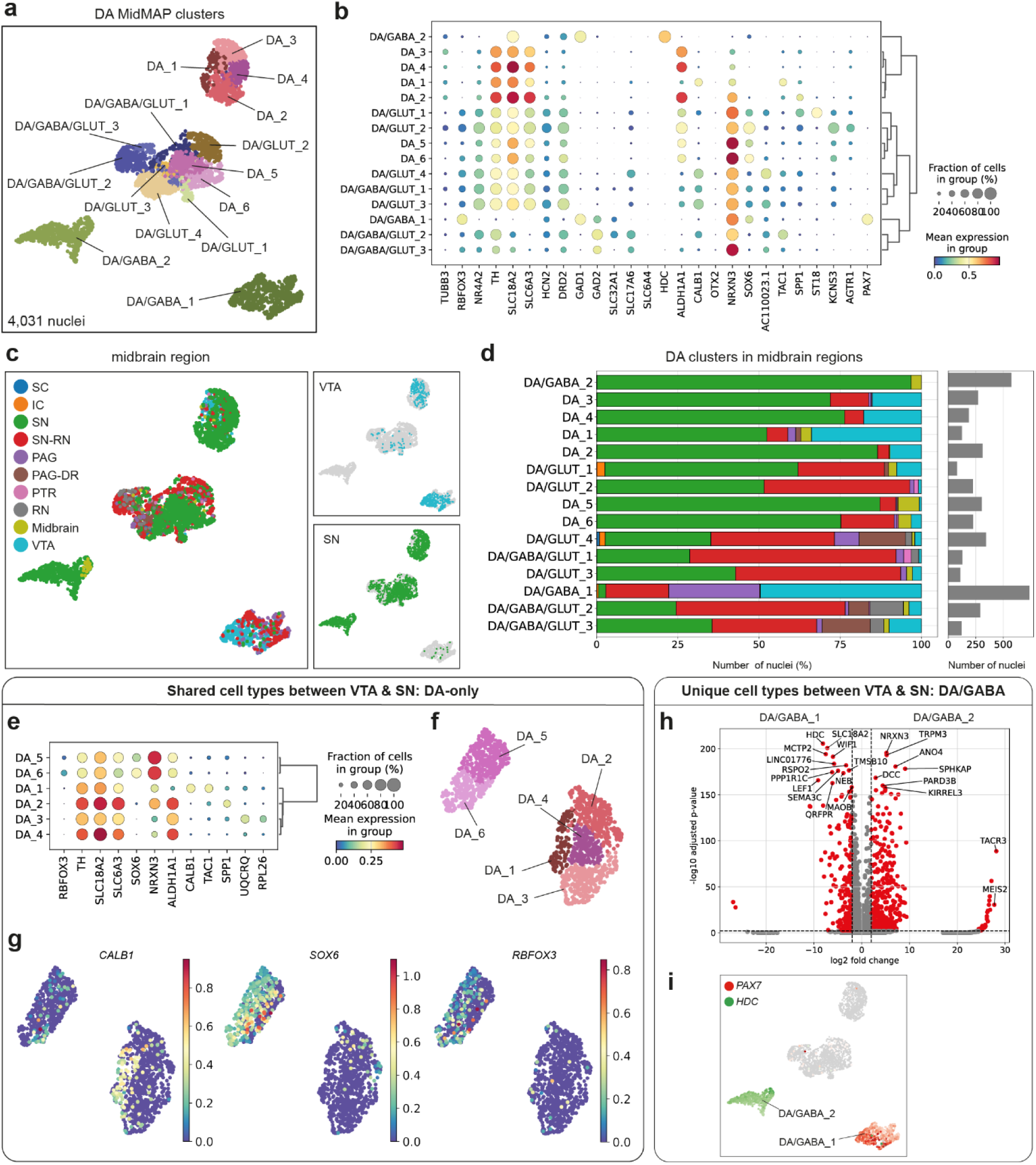
DA neuron diversity analysis reveals shared and unique cell types between midbrain regions. **a**, UMAP off the DA clusters in the integrated MidMAP atlas. **b**, Dot plot of marker gene expression in the DA clusters. **c**, UMAP of the midbrain regions, with the VTA and SN highlighted in separate boxes. **d**, Percentage of nuclei originating from the different midbrain regions, and the number of nuclei per cluster. **e**, Dot plot of marker genes in the DA-only clusters. **f**, UMAP showing the segregation of two main groups within the DA-only subclass. **g**, UMAPs with normalized gene expression of known region-specific genes. **h**, Volcano plot showing differentially expressed genes between the DA/GABA_1 and DA/GABA_2 clusters. **i**, UMAP showing cluster-specific gene expression of DA/GABA cell types within the DA neuron population.

**Fig. 6.**
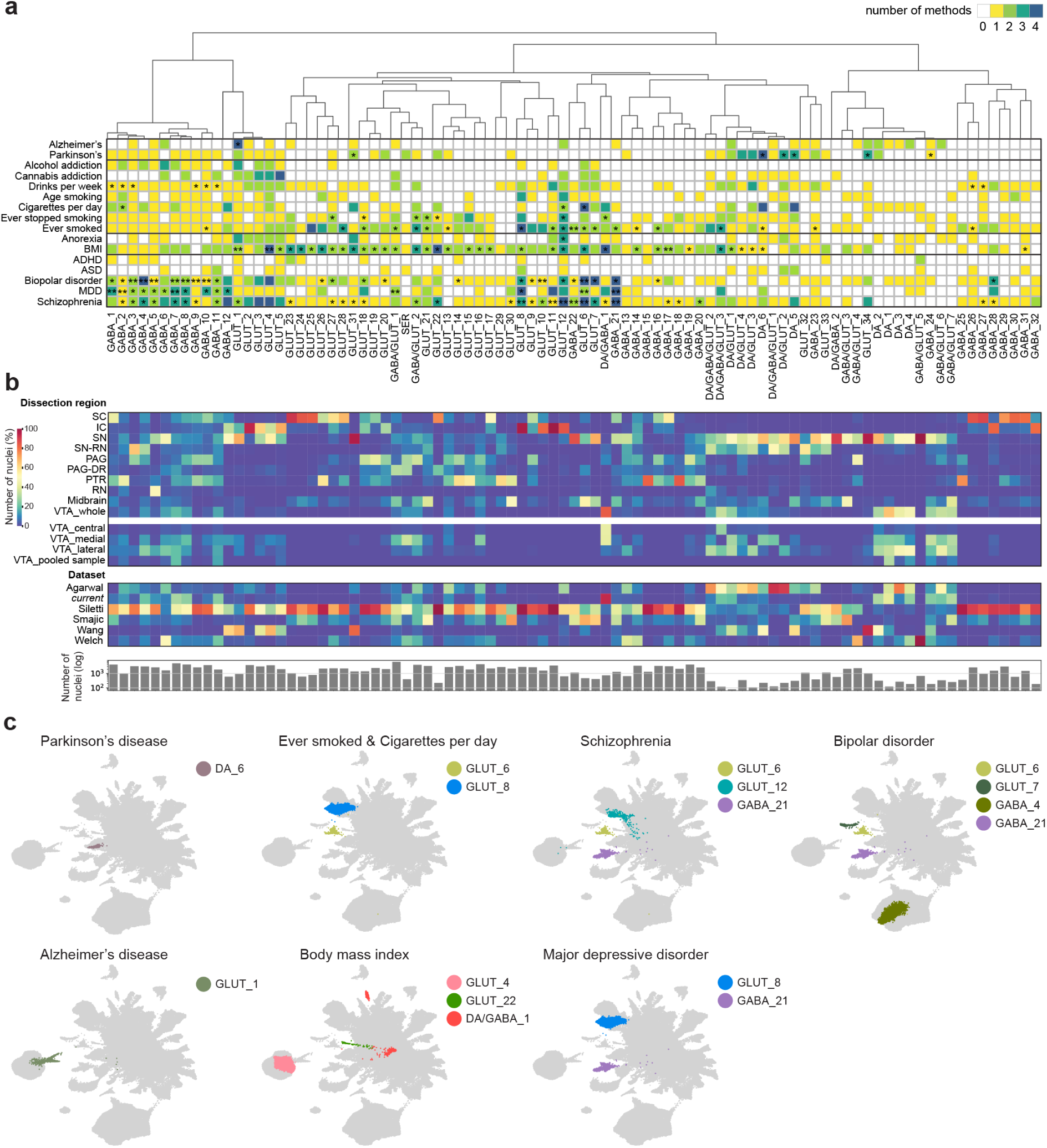
Trait-specific neuronal cell type identification in different brain disorders. **a**, Summarized plot of neuronal cell type associations in the human midbrain with different methods for multiple diseases and associated traits. Yellow boxes indicate a significant association (p=0.05) with one method, green with two, blue with three, and purple with four methods. Asterisks indicate the significance of the minimal p-value for the four methods used, after multiple testing correction for the number of cell types (*), and the number of cell types and traits (**). **Suppl.** Fig. 17 contains the results per method. BMI, body mass index; ADHD, Attention Deficit/Hyperactivity Disorder; ASD, Autism Spectrum Disorder; MDD, Major Depressive Disorder. SC, superior colliculus; IC, inferior colliculus; SN, substantia nigra; RN, red nucleus; PAG, periaqueductal grey; DR, dorsal raphe nucleus; PTR, pretectal region; VTA, ventral tegmental area. **b**, Percentages of the number of nuclei in the midbrain regions, specified per VTA subregion, and the total number of nuclei per cell type. **c**, UMAPs showing the cell types that have a significant association with all four methods that are also significant after multiple testing correction for the number of cells and traits.

We also used the integrated midbrain atlas to identify the VTA’s ‘unknown’ *FAT2*+ neurons. These nuclei clustered together with *SLC17A7*+ (VGLUT1) nuclei that also highly expressed *FAT2* and originated from the IC and SN **(Suppl. Fig. 12c, 12e).** Although the *FAT2*+ nuclei from the VTA did not express *SLC17A7*, these VTA nuclei may be GLUT neurons that we identify with our RNAscope experiments and may lack additional marker genes due to low sampling efficiency.

### DA neuron diversity in the human midbrain

Next, using the integrated human midbrain atlas, we investigated the DA neuronal landscape in more detail. We identified 4,031 DA neuronal nuclei that were grouped into 15 clusters (**Fig. 5a**, **Suppl. Fig. 15**). Annotation with known DA marker genes, such as *SOX6* and *CALB1*^24^, and DGE analysis revealed multiple subclasses and cell types (**Fig. 5b**, **Suppl. Fig. 16**). Most of the DA clusters originated from multiple midbrain regions, except for two DA-only and the DA/GABA clusters (**Fig. 5c**, **5d**). This revealed two features: DA clusters that are shared and that are unique among the VTA and SN (**Fig. 5e-i**), Within the DA-only subclass, DA_5 and DA_6 clustered together, differentially expressed *SOX6* and also *ALDH1A1*, and originated only from the SN, which confirms previous findings in rodents that identified this DA population in the SN (**Fig. 5c-g**)^3,36^. DA_1-4 originated from both the VTA and SN, indicating that both regions contain DA neurons with similar gene expression profiles. DA_1 showed high expression of *CALB1,* corresponding to the *CALB1*+ DA population described earlier^24^, whereas *ALDH1A1* was highly expressed in clusters DA_2, DA_3, and DA_4 **(Fig. 5e).** The two region-specific clusters in the DA/GABA subclass showed many differentially expressed genes, of which two were also unique within all DA clusters: *PAX7* in the VTA-specific DA/GABA_1 cluster, and *HDC* in the SN-specific DA/GABA_2 cluster (**Fig. 5b, 5h-i**). This indicates region-specific gene expression within the DA/GABA subclass that can also be used as marker genes for these neurons.

Together, our analysis identifies both VTA-specific cell types (DA/GABA) and four DA-only cell types that are present in multiple midbrain regions. Moreover, we reveal regional distribution and molecular signatures for human DA neurons in the whole midbrain, demonstrating the value of region-specific snRNA-seq datasets and their integration to identify region-specific cell types.

### Trait-specific neuronal cell type identification in different brain disorders

The VTA is associated with multiple psychiatric, developmental, and possibly also neurodegenerative disorders, but the contribution of specific cell types in the VTA remains unclear. To address this, we used Genome-Wide Association Studies (GWAS) summary statistics to link genetic risk for traits associated with the midbrain and VTA to gene expression profiles of the cell types that we identified in the human midbrain. Since there are many GWAS-to-cell type methods available that use different assumptions, we ran four methods (CELLECT^39^, FUMA cell typing^40^, scDRS^41^, and sc-linker^42^; **Methods**) on the integrated midbrain atlas to prioritize cell types and investigate whether they are DA cell type or brain region-specific.

To benchmark the analysis, we used Parkinson’s disease (PD) GWAS data. The integrated midbrain atlas includes *SOX6*+ DA cell types that are specific to the SN, which are the primary neuronal population affected in PD^66^. Accordingly, the two SN-specific *SOX6*+ DA-only cell types (DA_5 and DA_6) showed significant enrichment for PD (**Fig. 6a, Suppl. Fig. 15**)^43^. Additional associations with a DA/GLUT cell type (DA/GLUT_2) and two SN-originating *SLC17A7*+*FAT2*+ GLUT cell types (GLUT_31 and GLUT_34) suggests involvement of another SN-specific subclass, supporting studies on non-dopaminergic dysfunction in this disease^44–47^ (**Fig. 6a**, **6b**).

The dopamine system has also been linked to the psychiatric symptoms that arise in patients with Alzheimer’s disease (AD) such as apathy, depression, and psychosis^48^. One *SLC17A7*+*FAT2*+ GLUT cell type that originated from the SN/IC (GLUT_1) showed a significant association with AD, which implies a possible indirect effect of reward system dysfunction in the VTA in AD, or the involvement of glia^49–52^.

Substance-use disorders including alcohol addiction, cannabis addiction, and smoking have been associated with an altered dopaminergic output from the VTA in the reward system^53,54^. The traits “cigarettes per day”, “ever smoked” and “ever stopped smoking” showed multiple specific associations, including the triple combinatorial DA/GABA/GLUT_3 containing VTA-originating nuclei and SN-specific DA_6 for “ever smoked”, and the VTA-specific cell type DA/GABA_1 for “ever stopped smoking”. Cannabis addiction did not correlate to a specific cell type. Six *OTX2+* GABA cell types that also contained VTA nuclei (GABA_1-3 and GABA_9-11) were associated with “drinks per week”, indicating a possible role for VTA GABA neurons in this trait.

Eating disorder-related traits such as anorexia nervosa (AN) and body mass index (BMI) are also associated with the reward system^55^. AN showed one association with a GLUT cell type (GLUT_12) that originated exclusively from the IC. Notably, this cell type had the most associations with diverse traits, indicating that this cell type could be implicated in cross-disorder mechanisms. BMI had many strong cell type associations, suggesting that several cell types in the midbrain are involved in this trait, possibly reflecting the highly heterogeneous and polygenic etiology of this trait. Also, significantly associated cell types with this trait in the VTA originated from different VTA subregions, such as GLUT_4 mostly from the medial and lateral parts, and DA/GABA_1 mostly from the central and medial parts, suggesting heterogeneous cell type and regional association within the VTA as well.

Several developmental disorders have been associated with the reward system, including Attention Deficit/Hyperactivity Disorder (ADHD), Autism Spectrum Disorder (ASD), Bipolar Disorder (BP), Major Depressive Disorder (MDD), and schizophrenia (SZ). *OTX2*+ GABA cell types that originated from multiple midbrain regions were strongly associated with BP, MDD, and SZ, as well as a few GLUT neurons from different midbrain regions including the SN, SC, and IC. Notably, the majority of the VTA-originating *OTX2+* GABA cell types that were associated with these traits were sampled from the medial and lateral VTA, suggesting VTA subregion-specific implication. Also, the VTA-specific DA/GABA_1 cell type showed an association with SZ. ADHD and ASD did not show any significant cell type associations.

Overall, traits showed associations with cell types across midbrain regions. Among these were *OTX2*+ GABA cell types, also originating from the VTA, which were associated with neuropsychiatric traits, a VTA-specific DA/GABA cell type with BMI, and DA neurons with PD, BMI, and “ever smoked”. Some traits showed overlapping cell types that were identified by all four methods (**Fig. 6a, 6c**), suggesting shared genetic mechanisms and cell types in reward system-related traits. Together, these results implicate multiple cell types and brain regions in reward system-related brain disorders, and that non-DA midbrain neurons may be the primary candidates for many of these disorders.

## Discussion

To address a gap in knowledge of the neuronal cell type composition in the human VTA, we generated a transcriptomic reference atlas consisting of over 85,000 nuclei using state-of-the-art snRNA-seq. Using this unique dataset, we identified cell types specific to the VTA subregions even though most were distributed across the whole VTA. Moreover, we provide the first identification of functional connectivity differences between these VTA subregions. Integration of our VTA atlas with published human midbrain datasets allowed us to investigate the DA neuron population and identify multiple disease-associated neuronal cell types from GWAS summary statistics. Therefore, this study provides a comprehensive multimodal view of the adult human VTA.

### Neuronal diversity and connectivity-based organization in the adult human VTA

A major function of the VTA is its DA output, which is highly regulated by a local heterogeneous neuronal microcircuit^56–58^. The neuronal diversity we found in the human VTA is consistent with rodent studies, suggesting a conserved neuronal system^2,3,5,59^. Also in the human VTA, key determinants of neuronal populations were transcription factor, axon guidance and neuropeptide genes. These regulatory programs play a key role in regulating cell type identity and function, and can therefore provide entry points for targeted manipulation in experimental models^60^. Despite using the anatomical subdivision of the VTA into three regions, our findings suggest that its spatial organization is not defined by discrete, region-specific subclasses and cell types. Instead, regionalization probably relies on gene expression gradients, exemplified by *SOX6* expression in cell types located in the central VTA, and supported by the transcriptional similarity between VTA neurons and adjacent midbrain regions such as the SN. While certain VTA subregions show enriched neuronal populations, such as *OTX2*- GABA neurons in the central VTA or previously described DA/GLUT neurons in the medial VTA^14^, the overall distribution of neuronal subclasses remains largely homogeneous. Nevertheless, distinct cell types could have specific functions in local or broader connectivity networks, and future studies could focus on neuronal ensembles in VTA subregions. It is noteworthy that our pragmatic clustering strategy did not aim at delineating cell types using highly refined clustering, which results in the range of thousands. Their diversity could be analyzed in more detail as we did for the *TH*-high clusters in the integrated atlas to find regional differences.

Consistent with rodent studies emphasizing projection-defined neuronal organization, our data further suggests that neuronal localization within the VTA plays a key role in shaping connectivity, independent of neurotransmitter profile^9,21,61^. For example, the medial VTA is strongly associated with the somatomotor-related connectivity, which include the primary motor, sensory, and auditory cortices, and could correlate with the DA/GLUT neuron enrichment in this region. Such connections between sensory and motor cortices and the VTA are supported in humans with structural connectivity^63^ and in rodents with functional studies^64,65^, though the subregional organization within the VTA had not been investigated in humans yet. Together, these observations indicate that functional specialization arises from a combination of differences in cell type composition and regionally-driven connectivity profiles, though a true causal link can only be established with further functional studies.

### Dopaminergic neuron diversity in the adult human midbrain

We leveraged published single-cell studies of the human SN, which captured the DA cell type composition. By integrating these with our VTA-specific snRNA-seq dataset, we aimed to both refine VTA cell types and to compare VTA and SN populations. DA neurons are likely undersampled, and the higher number of DA nuclei allowed us to identify four DA-only clusters in the VTA instead of one in the VTA-only dataset. Similarly, Kamath et al. described 10 DA cell types in the human SN as opposed to 15 in our integrated midbrain atlas^28^. The addition of more donors, or single-nucleus or spatial RNA-seq datasets might therefore reveal further diversity^28^. Importantly, our findings expand the current view of the DA neuron organization within and between different regions of the human midbrain by showing that DA cell type identity is not strictly region-bound but instead spans both shared and region-specific populations across the VTA and SN. Through this, we move beyond prior regional bulk RNA-seq studies and provide the first detailed characterization of regional DA cell types in the human midbrain. While *CALB1* and *SOX6* remain distinguished marker genes between human DA neurons^24,28^, our data reveal additional subpopulations, including neurons expressing *ALDH1A1* – a marker previously identified in rodent studies^36^ - but lack *SOX6* or *CALB1* expression, highlighting greater molecular heterogeneity than previously described. Furthermore, we show that the DA/GLUT_1-3 cell types in the VTA correspond mostly to either the rodent *VGLUT2*+*ALDH1A1*+*OTX2*+ or the *VGLUT2*+*ALDH1L1*- cell type^8^, because they expressed *ALDH1L1+* but not *OTX2+*. Additionally, these cell types expressed *SOX6*+, which has not been described in combination with *VGLUT2* in the rodent midbrain. Therefore, comparison with rodent studies indicate only partial overlap in marker gene combinations and cell types highlighting that there is a need to identify human-specific molecular signatures and cell types in the VTA and SN^2,28^.

### Implicated cell types in reward system-related traits

Multiple brain disorders that have been linked to an imbalanced DA system have polygenic etiology, which makes the identification of implicated cell types challenging. Our polygenic risk analysis revealed a clear difference between DA and non-DA neurons, with a specific association of DA neurons with neurodegenerative disorders and other neuronal subclasses with neuropsychiatric traits. GABA neurons, increasingly recognized for their involvement in neuropsychiatric diseases, are key regulators of dopamine output from the VTA, suggesting that dysregulation of local inhibitory control may contribute to disease-related dysfunction^58^. Moreover, the presence of trait-associated cell types in multiple midbrain regions, such as the *OTX2*+ GABA_1-12 cell types, suggests that traits arise through both region-specific connectivity and local network mechanisms.

Correlating cell types to connectivity profiles across VTA subregions highlights how region-specific disease-associated neuronal populations align with functional brain networks. For example, the somatomotor network, previously linked to BMI^67^, showed the strongest connectivity with the medial VTA. Notably, the most significantly BMI-associated cell type, GLUT_4, was predominantly located in this region, suggesting a link between a VTA-located glutamatergic cell type and the somatomotor network in BMI. This reflects the involvement of VTA glutamatergic neurons in reward, feeding and locomotion^68^, but a direct projection of these neurons to the somatomotor cortex has not been described in the rodent brain. Additionally, GABA_1-11 and GABA_21, associated with BP, MDD, and SZ in our polygenic risk analysis, predominantly originated from the medial and lateral VTA. We show that these regions are connected to the default mode network and subcortical structures such as the hippocampus, which have extensively been associated with psychiatric disorders^69–72^. Together, our findings suggest that specific subregional VTA cell types contribute to disease-relevant brain networks, providing new insights into how specific circuits may underlie psychiatric disorders.

### Future perspectives

Knowing which neuronal cell types are associated with diseases, their gene expression profile, and in which brain region they are present is invaluable for studying these disorders in model organisms. One future aspect is the comprehension of the functional behavior of disease-implicated cell types. For this, the use of animal models will be necessary to study cell type-specific connectivity and disease-related behavior. A comparison of VTA snRNA-seq datasets of multiple species will be a crucial step in finding gene expression profiles of functionally similar cell types, which can be exploited for genetic and viral targeting for mechanistic studies. The human-specific disease-associated cell types that this study has revealed are potential targets for developing new treatment strategies for reward system-related diseases. Altogether, this study takes us a step further in understanding the molecular and cellular mechanisms underlying the reward system and its diseases.

## Methods

### Experimental methods

#### Human brain samples

Human post-mortem brain tissue from non-demented control donors was obtained from the Netherlands Brain Bank (NBB, https://www.brainbank.nl/) (**Suppl. Table 1, Suppl. Fig. 1a**). All donors gave informed consent to use their brain tissue for research after death. The NBB only reveals an autopsy serial number for donor anonymity, which includes the year and the number of the autopsy. Fresh frozen cryopreserved midbrain tissue (n=6) and formalin-fixed paraffin-embedded (FFPE) (n=3) were kept at –80 °C and room temperature (RT), respectively, until further processing. The NBB creates a short post-mortem delay by performing quick autopsies to ensure high tissue quality, which resulted in a mean post-mortem delay of around seven hours for the cryopreserved brain tissue.

The procedures of the NBB have been reviewed and agreed to by the medical ethics committee of the Amsterdam University Medical Center, the Netherlands (2009/148). These concern the donation of brain material by donors after their death for scientific purposes. The scientific committee of the NBB has reviewed and approved this study. The NBB follows BrainNet Europe’s Code of Conduct for brain banking in obtaining and handling human brain tissue, which includes ensuring quality, safety, and ethics in all procedures. The relevant regulations that involve the use of human participants were followed by this study, as described by the Declaration of Helsinki and the Dutch and European legal and ethical criteria.

#### Nuclei isolation of human post-mortem VTA tissue

Fresh frozen post-mortem human midbrain tissues were cryosectioned on a dorsal-ventral axis into 250 µm thick sections (**Suppl. Fig. 1a-c**). The VTA was identified by the presence of the red nucleus and the pigmented substantia nigra. Per section, the medial, central, and lateral subregions of the VTA were manually dissected with a cooled scalpel and collected in pre-cooled Eppendorf tubes on dry ice (**Fig. 1a**, **Suppl. Table 2**). In three donors the tissue blocks only contained one VTA subregion due to the cutting angle during the autopsy, and from one donor the three regions were pooled due to low nuclei counts. Nuclei were isolated as previously described with a few adjustments^28,73,74^. In short, all procedures were performed on ice and in a fume hood to avoid dust particles. Cryosectioned VTA tissue was transferred to a pre-cooled 2 mL KIMBLE Dounce tissue grinder set (D8938, Sigma-Aldrich), to which 1 mL Lysis Buffer (10 mM Tris, 10 mM NaCl, 3 mM MgCl_2_, 10 mM CaCl_2_, 0.1% NP40 in nuclease-free water (NFW; W4502, Sigma-Aldrich)) was added. The tissue was homogenized using 5 times Pestle A and 15 times Pestle B included in the douncer set and transferred to a new pre-cooled 50 mL tube. The douncer was rinsed with 4 mL Lysis buffer, which was added to the 50 mL tube and incubated on ice for 15 minutes with gentle mixing during the incubation. After the incubation, 5 mL Hybernate A medium (A1247501, Thermo Fischer Scientific) containing 0.01% GlutaMAX (35050061, Thermo Scientific) was added to the tissue solution and resuspended five times with a 5 mL pipette. The solution was filtered using a 40 µm pluriStrainer cell strainer (43-50040-51, pluriSelect) into a new pre-cooled 50 mL tube and spun down in a swing-bucket centrifuge at 500 x g at 2 °C for five minutes. 1 mL Nuclei Wash Buffer (1X PBS, 2% BSA, 0.2 U/µL Protector RNAse Inhibitor (3335402001, Sigma-Aldrich), 0.22 µm-filtered before use) was added to the pellet, resuspended, transferred to a new pre-cooled 1.5 mL Eppendorf tube and centrifuged at 500 x g at 2 °C for five minutes. The pellet was resuspended in 500 µL Sucrose Nuclei Solution (946 µL Nuclei Wash Buffer + 154 µL 1.8 M Sucrose Solution; 1.8 M sucrose, 10 mM Tris, 10 mM NaCl, 3 mM MgCl_2_, 10 mM CaCl_2_, 0.2 U/µL Protector RNAse Inhibitor in NFW) and carefully pipetted on top of 500 µL 1.8 M Sucrose Solution in a new 1.5 mL Eppendorf tube. After centrifugation at 13.000 x g at 4 °C for 45 minutes, the supernatant containing myelin and cellular debris was carefully removed. The pellet was washed once by adding 1 mL Nuclei Wash Buffer to the pellet, gently resuspended, and centrifuged at 500 x g at 2 °C for five minutes. The pellet was resuspended in 500 µL to 1000 µL Nuclei Wash Buffer depending on the amount of tissue at the start of the protocol, and filtered using a 40 µm cell strainer into a new 1.5 mL Eppendorf tube. 5 µL was taken from each sample to confirm good cell lysis using a Countess Automated Cell Counter (Thermo Fischer Scientific) using Trypan Blue with a cell viability of around 5% as a good indication of cell lysis (10X Genomics). DAPI (1:200, D9564, Sigma-Aldrich) was added to the nuclei solution and incubated on a rotator at 4 °C for 15 to 30 minutes, after which the solution was filtered using a 40 µm cell strainer in a BSA-coated sterile fluorescence-activated cell sorting (FACS) tube (3% BSA, 1X PBS; incubation at RT). The nuclei were sorted with a BD FACSAria II cell sorter (Becton Dickinson Biosciences) and BD FACSDiva software (BD Biosciences). Single nuclei were selected based on size and DAPI signal, sorted with an 85 µm nozzle, collected in 3% BSA-coated 1.5 mL Eppendorf tubes, and kept on ice until further processing (**Suppl. Fig. 1e**). Nuclei were visually inspected under a widefield microscope before and after fluorescence-activated nuclei sorting (FANS) to confirm the presence of intact nuclei (**Suppl. Fig. 1f**).

#### Droplet-based single-nucleus RNA sequencing

12 nuclei samples were collected and prepared for RNA sequencing by Single Cell Discoveries, The Netherlands (**Suppl. Table 2**). Per sample, 10,000 nuclei were loaded on a Chromium single cell 3’ Chip for v3.1 10x technology using the 10x Genomics V3 kit. Single nuclei were encapsulated into gel-made droplets together with a barcode, and lysis and reverse transcriptase reagents. Reverse transcription of barcoded RNA was followed by cDNA amplification, and libraries were made from each sample individually. Libraries were loaded and sequenced on Ilumina NovaSeq 6000 with a target of 40,000 reads per nucleus.

#### Immunohistochemistry

FFPE human midbrain tissue from non-demented control donors was cut into 7 µm thick sections using a microtome (RM2155, Leica), collected on Superfrost plus slides, and dried in an oven at 37 °C overnight. The section was deparaffinized and rehydrated by submersion in xylene two times for 10 minutes each, a series of decreasing ethanol dilutions (two times 100%, 96%, 80% 70%, and 50%: two minutes each), and finally in demi water. After washing the section in 1X PBS/0.05% Tween for 10 minutes, antigen retrieval was performed with pre-warmed 10 mM citrate buffer (pH 6.0) supplemented with 0.05% Tween in a steamer for 20 minutes. After cooling at RT for 20 minutes, the section was washed in 1X PBS/0.05% Tween and blocked with blocking buffer 1X PBS/1% NHS/0.1% BSA/0.2% TritonX100 at RT for one hour (NHS, Normal Horse Serum, ab7484, Abcam; BSA, Bovine Serum Albumin, 9048-46-8, Sigma-Aldrich; TritonX100, 9036-19-5, Sigma-Aldrich). The section was incubated with rabbit-anti-TH primary antibody (1:500, AB152, Millipore) in blocking buffer at 4 °C overnight, washed in 1X PBS/0.05% Tween two times 5 minutes each, and incubated with fluorescently labeled secondary antibody donkey-anti-rabbit-AF568 (1:1000, ab175470, Abcam) in blocking buffer at RT for one hour. After three washes in 1X PBS/0.05% Tween for 10 minutes each, autofluorescence was quenched by incubating with True Black (23007, Biotium, lot: 22T0331) for 30 seconds, followed by three washes in 1X PBS for five minutes each. Nuclear staining was performed by incubation with Hoechst (1:1000 in 1X PBS/0.1% BSA, Invitrogen 33258, lot: 10778843, Thermo Fischer Scientific) at RT for 10 minutes. The section was washed in 1X PBS two times three minutes each, mounted in FluorSave (Merck Millipore, lot: 3861305), and stored at 4 °C.

#### In situ hybridization with RNAscope

In situ hybridization was performed on a 7 µm tick FFPE tissue section from the midbrain of three non-demented control donors (**Suppl. Table 1**) with an RNAscope kit (323180, 310091, 323135, ACD Biotechne, Newark, Canada), which was used according to the manufacturer’s instructions. The section was cut, mounted on Superfrost plus slides, and dried at 37 °C overnight. Deparaffinization of the section was performed with xylene two times for 10 minutes each, and a decreasing ethanol gradient (two times 100%, 96%, 80%, 70%, and 50% for two minutes each) followed by demi water. The section was incubated with Hydrogen Peroxide solution provided by RNAscope to block endogenous peroxide at RT for 10 minutes. After a brief wash in demi water, the section was incubated in pre-warmed 1X Target Retrieval Reagent in a steamer for 20 minutes and cooled at RT for another 20 minutes. RNAscope Protease Plus was added to the section and incubated at 40 °C in a HybEZ oven within the HybEZ Humidity Control Tray for 15 minutes. After a two-times wash in demiwater with mild rocking, pre-warmed probes for Hs-*TH* (441651, Biotechne), Hs-*GAD1* (404031, Biotechne), and Hs-*SLC17A6* (415671, Biotechne), were added to the section and incubated in the HybEZ oven at 40 °C for two hours. The section was washed in RNAscope 1X Wash Buffer, incubated with RNAscope Multiplex FL v2 Amp1 at 40 °C for 30 minutes, RNAscope Multiplex FL v2 Amp2 at 40 °C for 30 minutes, and RNAscope Multiplex FL v2 Amp3 at 40 °C for 15 minutes, with wash steps with 1X Wash Buffer of two times two minutes each in between the incubations. First, the *GAD1* probe signal was developed by incubation of RNAscope Multiplex FL v2 HRP-C2 at 40 °C for 15 minutes, washing twice with 1X Wash Buffer for two minutes each, and incubation with TSA vivid Fluorophore 570 (1:1500, diluted in RNAscope TSA dilution solution) (7526, Tocris ACD) at 40 °C for 30 minutes. The section was washed twice with 1X Wash Buffer for two minutes each, incubated with RNAscope HRP blocker at 40 °C for 15 minutes, followed by two washes in 1X Wash Buffer for two minutes each before the signal development of the next probe, *SLC17A6*. The sections were incubated with RNAscope Multiplex FL v2 HRP-C3 at 40 °C for 15 minutes, washed in X Wash Buffer two times two minutes each, and incubated with TSA vivid Fluorophore 650 (1:1500, diluted in RNAscope TSA dilution solution) (7527/2, Tocris ACD) at 40 °C for 30 minutes. To avoid the autofluorescence signal of human brain tissue detected around 488 nm, the third probe signal was developed in the near-invisible far-red spectrum, for which we used the Opal 780 reagent pack (FP1501001KT, Akoya Biosciences). First, the section was incubated with RNAscope Multiplex FL v2 HRP-C1 at 40 °C for 30 minutes and washed in RNAscope 1X Wash Buffer twice for two minutes each. Then, TSA-DIG (1:100, diluted in RNAscope TSA dilution solution) was added and incubated at RT for 30 minutes and washed in RNAscope 1X Wash Buffer twice for two minutes each. RNAscope Multiplex FL v2 HRP-C1 was added to the section and incubated at 40 °C for 15 minutes, washed twice in RNAscope 1X Wash Buffer for two minutes each, and incubated with Opal 780 Reagent (1:25, diluted in TBS-0.1% BSA; 1X TBS, Tris Base, NaCl, pH 7.6) at RT for 30 minutes. The section was washed in RNAscope 1X Wash Buffer twice for two minutes each, counterstained with RNAscope DAPI, which was incubated at RT for 30 seconds, briefly washed in 1X PBS, mounted in FluorSave (Merck Millipore, lot: 3861305), and stored at 4 °C.

#### Imaging and analysis

Imaging of the TH immunohistochemistry staining was performed on the DMi8 widefield microscope (Leica) with a 10x/0.32NA objective (11506521, Leica). The fluorescent signal was visualized at 568 nm. Imaging of the RNAscope staining was performed on a Leica RMI THUNDER Widefield Microscope (inventory number 359612) with a 20x/0.4NA objective (HC PL FLUOTAR L 20x/0.40 CORR PH1, 11506243, Leica) or 20x/0.8NA objective (HC PL APO 20x/0.8, 11506529, Leica), and LED8 light source (11504256, Leica). Fluorescent signal was visualized at 405 nm, 568 nm, and 647 nm using filter block DFT51010 (11525418, Leica), and 750 nm using filter block CYR71010 (11525416, Leica), and captured with a Leica K8 Camera (11547116) with a pixel resolution of 2048 x 2048. A single-plane image of the whole section was made with Leica Application Suite (LAS) X Navigator software, where a Focus Map was made to interpolate the z position. Z-stacks of several individual neurons were acquired on the STELLARIS 8 Confocal Microscope (Leica) with a 40x/1.3NA objective, a step size of 0.5 µm, and pixel resolution 512 x 512 with a voxel size of 0.284 x 0.284 x 0.346 µm. Fluorescent signal was visualized with a 405 nm laser, and White Light Laser with a tunable range of 440-790 nm. Multispectral imaging was performed using detectors HyD S for positions 1 and 3, HyD X for positions 2 and 4, and HyD R for position 5. Identification of probe signals in neurons and thus neuronal subclasses was based on the higher number of probe signal punctea in the nucleus and/or cell body of a cell compared to other cells and the background signal in the same section, as indicated by RNAscope. Maximum intensity projections of the Z-stacks were made with Imaris software (version 10.1.1). Individual neurons were located manually with ImageJ software (version 1.54f), and their coordinates were plotted using RStudio version 4.3.3.

### Computational methods

#### snRNA-seq data processing

##### Adult human VTA atlas generation

The raw count data of each of the 12 samples were aligned to the human reference genome GRCh38 (GENCODE v32/Ensembl98) using Cell Ranger (version 3.1.0) with the ‘--include-introns’ parameter. We used Scanpy (version 1.9.6) to perform our snRNA-seq analysis. Quality control of individual samples included filtering out ambient RNA using CellBender^75^, and low-quality nuclei based on the number of unique molecular identifiers (UMIs) with a cut-off range between 1000 and 10,000 counts, doublets based on a calculated doublet score with a cut-off of <0.2, <3% mitochondrial genes, <25% ribosomal genes, and a minimum number of cells of 3. This resulted in 92,937 nuclei in total, with a median of 3,064.8 genes and 8,355.9 UMIs per cell (**Suppl. Table 2**). We identified the highly variable genes and excluded *MALAT1* because it was highly expressed in all samples and not informative for cell type-specific marker gene discovery. All the samples were checked for *TH* expression as a quality measure of isolating DA neurons and subsequent sample inclusion in the dataset. The samples were integrated using Harmony using the Seurat pipeline to remove batch effects for the day of sample processing (10x batch) and donor^76^. Dimensionality reduction was performed using principal component analysis (PCA), and the neighborhood graph was calculated using 50 principal components (PCs), and UMAP with a minimum distance between embedded points of 0.5. Grouping of nuclei into clusters was initially computed by the Leiden algorithm to identify cell classes and for quality control in pre-processing steps. The final clusters were computed on the whole dataset using random walks with the Walktrap algorithm using the cluster_walktrap() function of the igraph package, based on the SNN neighborhood graph generated using the BuildSNNGraph() function of the scran package, which resulted in 93 clusters with a modularity score of 0.701^77^. Based on gene expression of known marker genes of cell classes, clusters with high expression of two or more marker genes for different cell classes (e.g. clusters with expression of astrocyte marker *GFAP* and also microglia marker *AIF1*) were identified as doublets and removed from further analysis, after which 85,254 nuclei grouped in 68 clusters remained for downstream analysis. Highly variable genes, PCA, and UMAP were recalculated to identify cell classes based on known marker gene expression for neurons, glia, and other non-neuronal cell types, and their hierarchical relationships. The neuronal cell class was isolated and its highly variable genes, PCA, and UMAP recalculated. Neuronal subclasses were identified based on known neurotransmitter-based gene expression. Differential gene expression analysis was performed between the neuronal clusters using a Wilcoxon rank-sum (Mann-Whitney-U) test to identify marker genes. Within neuronal subclasses (e.g. GABA), clusters that expressed similar marker genes were grouped to call cell types (e.g. GABA_SEMA5A/SST).

##### Adult human midbrain atlas generation

###### Data collection

**Suppl. Table 3** contains an overview of all atlases included in the integration and their metadata. For two atlases, Smajic et al.^27^ and Agarwal et al.^23^, the fastq files were downloaded using fastq-dump and donor SRR identifiers. To access the fastq files from Welch et al.^26^, the 10X BAM files were downloaded directly from the Sequence Read Archive and subsequently converted to fastq files with the dedicated 10X bamtofastq function. Fastq files from Wang et al. were downloaded through the dedicated Human Cell Atlas Data Explorer and filtered on donors. Lastly, midbrain fastq files from Siletti et al.^24^ were manually selected and downloaded from the Neuroscience Multi-Omic (NEMO) Archive. We used the metadata provided by Siletti et al. to assign cells to a specific region. Other datasets were derived entirely from a single region or the entire midbrain.

##### DA neuron cell type identification

###### Realignment to same reference genome

From the fastq files, alignments were performed with Cell Ranger v6.1 using the Ensemble 109 human reference genome, including the ‘--include-introns’ parameter. Additionally, the ‘--expect-cells’ parameter was set to the number of cells found in the samples of the original studies. Cell Ranger’s unfiltered count tables were used to remove ambient RNA and score the probability of droplets containing a cell using Cellbender 2.0 over 150 epochs^75^. The filtered output of Cell Ranger was used to estimate the average number of cells per sample and the number of background droplets.

###### Quality control

Cellbender’s filtered count tables were preprocessed with Scanpy (version 1.9.6). For each atlas, we annotated the cells with metadata features including species, atlas, and sample-id. Quality control metrics were added with the calculate qc metrics function. Cells with fewer than 500 reads were filtered out automatically through Cell Ranger. Mitochondrial genes and genes with fewer than three reads were filtered out, as well as cells with more than 10% mitochondrial or ribosomal reads. Doublets were detected and filtered using scrublet. Expected doublet rates were manually calculated with the guidelines from 10X Genomics based on the number of recovered cells in the original studies^78^.

Normalization was performed on cell-level with ‘normalize total’ using a total sum of 10000. Subsequently, log1p was used to condense the range of values and reduce the influence of highly expressed genes. After adding the original cell type annotations, the datasets of each study were combined as input for the integration process.

##### Integration of six snRNAseq datasets

Following preliminary integrations with scVI, Harmony, and Scanorama. The final integration was performed with scVI using 2000 highly variable genes, 90 latent features, 5 hidden layers, and 128 hidden nodes.

###### Label harmonization

Original annotations from each dataset were divided into three levels, cell type lvl1-3, one being the lowest resolution and three being the highest. Cells with fewer than three levels of annotation were represented by their highest resolution label in the higher level. Cell type hierarchy tree formation was performed using scHPL’s train tree function with the lowest resolution labels and the integrated latent space. The cell type labels from the Welch et al. and Smajic et al. datasets required further reduction to a lower resolution label. Cell type annotations were then collapsed to the first level labels of the hierarchy tree, forming a harmonized low-resolution annotation. Realignment of the original datasets revealed novel unannotated cells. Label transfer of the harmonized labels was performed using a k-nearest neighbour classifier with k=30 and uncertainty threshold of 0.2 on the whole dataset. The resulting (re)classified neurons were split off for further analyses.

157,244 predicted neurons were isolated from the integrated human midbrain atlas, which contained nuclei from all six snRNA-seq datasets. The 5,000 highest-expressing genes were used for further downstream analysis. After recalculation of highly variable genes, PCA, nearest neighborhood distance using 90 PCs, and UMAP with a minimum distance between embedded, nuclei were grouped using Leiden clustering (resolution 3) which resulted in 75 clusters. Neuronal subclasses were identified in the same manner as for the VTA, namely based on the expression of genes necessary for different neurotransmitter machineries in the identified clusters. Partition-based graph abstraction (PAGA) analysis^38^ was performed in Scanpy, and low-connectivity edges were trimmed by setting their threshold to 0.5.

##### DA neuron cell type identification

The integration of snRNA-seq datasets identified 52 additional DA-only neuronal nuclei from the VTA (155 total), due to the use of the later version of the reference genome and preprocessing steps tailored to the integrated midbrain atlas generation. To identify DA cell types, two clusters (clusters 41 and 42) that showed high expression of DA subclass markers *TH*, *SLC18A2* and *SLC6A3* were isolated and re-clustered with the Leiden algorithm (resolution 1), which resulted in 16 clusters, among which were six DA-only clusters (**Suppl. Fig. 13**). These clusters replaced clusters 41 and 42 in the neuronal midbrain integrated dataset, resulting in 89 clusters total that were used for further analysis. Differential gene expression of DA clusters was computed with a Wilcoxon rank-sum (Mann-Whitney-U) test.

#### Trait-specific cell type identification using GWAS summary statistics

##### Selection and preprocessing of GWAS data

We downloaded publicly available summary statistics for Alzheimer’s disease (AD)^79^, Attention-deficit/hyperactivity disorder (ADHD)^80^, Alcohol Use Disorder^81^, Anorexia Nervosa (AN)^82^, Autism Spectrum Disorder (ASD)^83^, Bipolar Disorder (BP)^84^, Body Mass Index (BMI)^85^, Cannabis Use Disorder^86^, Major Depressive Disorder (MDD)^87^, Parkinson’s Disease (PD)^88^, Schizophrenia (SZ)^89^, and several addiction-related phenotypes: age of first smoking cigarettes per day, drinks per week, ever smoking and smoking cessation^90^.

##### GWAS-to-cell-type methods

Methods that aim to use GWAS signals to prioritize cell types have different underlying assumptions and implementations. Below, we briefly describe three key components that create differences between these methods and their results. The first component is the approach to bridge the gap between GWAS data (collected at the SNP level) and the snRNA-seq data (collected at the gene level). The second component is the selection of genes that characterize cell types. This is done by selecting genes that are specific to a cell type, or by characterizing a cell type by the average expression of genes. The third component is the statistical test that formally tests for association of GWAS signal and cell type. For most methods, these statistical tests are implemented through either the MAGMA pipeline version 1.10^92^ or the LDSC regression framework^91^. We applied four different methods to prioritize cell types (FUMA cell typing^40^, CELLECT^39^, scDRS^41^ and sclinker^42^, details in **Suppl. Note 2**).

##### Multiple testing correction

For all traits, cell types, and methods, we report nominal p-values, values corrected for the number of tested cell types (89; alpha level of 0.00056), and values corrected for all cell types and traits tested (89*16; alpha level of 3.51x 10^-5).

### Neuroimaging data analysis

#### Participants and Magnetic Resonance Imaging

Data from the Human Connectome Project were used for this project^31^. The relevant data packages are part of the WU-Minn Human Connectome Project Data-1200 Subjects dataset. Subjects who had 7T resting-state functional magnetic resonance imaging (fMRI) available were selected. The initial dataset comprises data from 184 individuals across four different sessions. In this project, 7T imaging in the Resting State fMRI FIX-Denoised (Extended) package and the Structural Preprocessed for 7T (1.6mm/59k mesh) were used. Subjects with known quality control issues were removed leaving a total of 181 subjects for subsequent analysis. The data were acquired and minimally preprocessed by the Human Connectome Project. The preprocessing and acquisition protocols are described in detail elsewhere^95,96^. Additional preprocessing and a full overview of the methods for fMRI analyses can be found in **Suppl. Note 1** and **Suppl. Figure 9**.

#### Functional Connectivity Analysis

##### Seed-based VTA functional connectivity

For these analyses, we defined three seed areas within the left VTA. The base for the definition of those is a probabilistic 7T atlas of the VTA, defined by Trutti et al^97^. We binarized the atlas using a threshold of 0.1 and manually segmented it into three disjoint subregions - central, medial, and lateral - to match the dissected areas (**Suppl. Fig 6**). The average time series for each of the three seeds were extracted using fslmeants on each of the subregional masks on each preprocessed image^98^. After calculating and removing the regressors from the time series, the extracted time series from each VTA seed was included in a separate regression model to create an individual functional connectivity map for each subregion of the VTA per subject session using FSL FEAT^99^. Then, each of the four connectivity maps was averaged per subject and a group map was estimated from the 181 individual average functional connectivity maps using a sign-flipping permutation method under a threshold-free cluster enhancement - a family-wise error multiple testing correction method - for significance filtering^100,101^.

##### Sub-regional cortical activation map comparison

The cortex of each subject was parcellated into the Desikan-Killiany Atlas (aparc) and subcortical regions were segmented according to the aseg atlas^28,29^. The subject-specific anatomical parcellations were registered to the standard MNI152 space. Using these cortical and subcortical atlases, the average functional connectivity value to each of the VTA seeds by averaging all voxels per brain area for each of the three functional connectivity maps. To test for differential activation in VTA subregions, a linear mixed-effects model was fit per pair of seeds allowing for a random intercept per subject and per target brain area to account for the multi-level structure of data using the lme4 (v1.1) and lmerTest (v3.1) R packages (v4.2.0). The interaction between the effect of the seed VTA subregion and the target cortical/subcortical region was studied by including an interaction term in the model. The threshold for significance was adjusted to correct for multiple comparisons, factoring in the number of seeds (3), cortical regions (44) in the left hemisphere, and interactions (alpha = 2.8·10^-4^). Age, sex, and intracranial volume were included as covariates. The emmeans package (v1.11.1) was then used to obtain the estimated marginal means of the pairwise differences between the connection strength between each seed for each target region. The heatmap containing the contrasts in the estimated marginal means was created in Morpheus (Broad Institute). Five different nesting structures were compared and the model with the lowest Bayesian Information Criterion (BIC) was used (random intercept per subject; random intercept per subject and per target; random intercept per subject and per seed; and random intercept per subject, per seed and per target (nested and non-nested)). The exact formula used was:

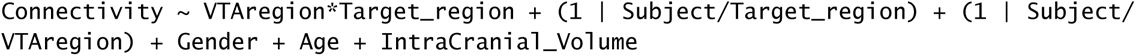

Besides the (sub)cortical areas of interest, the corpus callosum was included as a negative control: white matter regions should not have any specific activations (p_central-medial_ = 0.98; p_central-lateral_ = 0.98; p_lateral-medial_ =0.99). These analyses were repeated for the Von Economo-Koskinas Atlas for replicability purposes (**Suppl. Note 1**)^88^.

## Data availability

The snRNA-seq data from the adult human VTA are available for download on request.

## Code availability

The code used for analysis and the expression matrix of the adult human VTA snRNA-seq dataset will be available on Github.

## Acknowledgments

We thank Frank Meye and Kimberly Siletti for reading the manuscript, and Jacqueline Sluijs, Roland van Dijk, Keith Garner, and Youri Adolfs for technical support. We thank the Netherlands Brain Bank for contributing the human post-mortem brain tissue. We thank Single Cell Discoveries for processing the human VTA nuclei for snRNA-seq. We thank the Princess Máxima Imaging Center for microscopy support. This project was financially supported by the NWO Gravitation program BRAINSCAPES: A Roadmap from Neurogenetics to Neurobiology (NWO: 024.004.012) to E.M.H., O.B., M.P.H, R.J.P.

## Author contributions

A.S.v.R.A. conceptualized the study, designed all experiments, collected the human VTA nuclei for snRNA-seq, performed all snRNA-seq data analysis, performed and analyzed the RNAscope experiments, analyzed the fMRI and trait-specific cell type data, and wrote the manuscript. B.d.A.P.C.M. designed, performed and analyzed the fMRI experiments and wrote the manuscript. R.M.B. designed, performed and analyzed the trait-specific cell type analysis, and wrote the manuscript. C.J.R.J. integrated the human midbrain single-nuclei datasets and wrote the manuscript. S.L.S. performed the fMRI experiments. R.L.v.I. performed the confocal imaging of the RNAscope. J.M.E. performed and optimized the RNAscope experiments. T.M.H. helped optimizing the nuclei isolation protocol. A.C.R. provided the use of the Princes Máxima Center Imaging Facility. R.J.P. provided help in shaping the project and wrote the manuscript. V.D. conceptualized the study, supervised the project, and wrote the manuscript. M.P.v.d.H. provided support and feedback on the fMRI experiments. O.B. conceptualized the study, performed the human VTA snRNA-seq preprocessing and human midbrain datasets integration, supervised the project, and wrote the manuscript. E.M.H. conceptualized the study, supervised the project, and wrote the manuscript. All authors provided feedback during the project, and read and edited the manuscript.

