## Supplementary Materials for "Regional specificity of neuronal cell types, functional connectivity, and cell type-specific correlation to brain disorders in the human ventral tegmental area"

Contents

Supplementary Tables S1-S3
Supplementary Figures S1-S17
Supplementary Notes N1-N2

**Supplementary Table 1: Donor information of human midbrain tissue.**

| NBB number | sex | age | braak score | amyloid score | braaklb | pmd | pH | brain weight (gr) | ApoE |
|---|---|---|---|---|---|---|---|---|---|
| Fresh-frozen cryopreserved human midbrain tissue for snRNA-seq of the VTA | | | | | | | | | |
| 1996-052 | m | 73 | 2 | n.d. | 0 | 09:10 | ? | 1500 | 43 |
| 2007-075 | f | 82 | 2 | O | 2 | 05:10 | 6.64 | 1171 | 32 |
| 2007-082 | m | 81 | 2 | O | 0 | 07:55 | 6.23 | 1194 | 33 |
| 2008-027 | f | 80 | 1 | A | 2 | 06:55 | 6.50 | 1220 | 43 |
| 2008-103 | m | 80 | 1 | A | 1 | 08:10 | 6.18 | 1260 | n.d. |
| 2009-005 | m | 82 | 1 | O | n.d. | 05:10 | 6.75 | 1087 | 32 |
| Paraffin-embedded formalin-fixed midbrain tissue for immunohistochemistry and *in situ* hybridization | | | | | | | | | |
| 1998-104 | f | 74 | 2 | O | 0 | 07:25 | 6.95 | 1167 | 32 |
| 1999-101 | m | 69 | 1 | O | n.d. | 19:15 | 6.40 | 1337 | 33 |
| 2000-036 | m | 53 | 0 | O | n.d. | 14:2 | 6.58 | 1331 | 33 |

NBB, Netherlands Brain Bank; braaklb, braak lewy body; pmd, post-mortem delay; ApoE, apolipoprotein E genotype; n.d., not defined.

**Supplementary Table 2: Sample information and quality control (QC) metrics for snRNA-seq.**

| Sample ID | Donor ID | Donor number | VTA region | Nuclei number | Nuclei number after QC | Median genes per nucleus | Median reads per nucleus |
|---|---|---|---|---|---|---|---|
| g004 | 2007-075 | 1 | Central | 6771 | 6466 | 3673.5 | 11270.0 |
| g005 | | | Medial | 8811 | 8296 | 3307.5 | 9168.0 |
| g011 | | | Lateral | 9259 | 8808 | 3060.0 | 8095.0 |
| g012 | 2008-103 | 2 | Central | 2880 | 2721 | 3515.0 | 10657.0 |
| g013 | | | Medial | 10877 | 10229 | 2776.0 | 7034.0 |
| g014 | | | Lateral | 7338 | 6502 | 2642.0 | 6407.0 |
| g015 | 2007-082 | 3 | Lateral | 7712 | 6956 | 2743.5 | 6877.5 |
| g016 | 2008-027 | 4 | Lateral | 2323 | 2212 | 3194.5 | 10221.5 |
| g017 | 2009-005 | 5 | Pooled sample | 9630 | 8886 | 3014.5 | 7407.0 |
| g018 | 1996-052 | 6 | Lateral | 8834 | 8214 | 3139.0 | 8366.0 |
| g019 | | | Medial | 9235 | 8129 | 2963.0 | 7259.0 |
| g020 | | | Central | 9310 | 7835 | 3233.0 | 8632.0 |

**Supplementary Table 3: Midbrain dataset integration and quality control metrics.**

| Atlas | Midbrain region | Nuclei number | Median genes per nucleus | Median reads per nucleus |
|---|---|---|---|---|
| Agarwal et al. (2021) | SN | 8178 | 930 | 1325.5 |
| van Regteren Altena et al. (current) | VTA | 84503 | 2978 | 7716 |
| Siletti et al. (2023) | Multiple midbrain regions | 282660 | 3200 | 7682.5 |
| Smajic et al. (2021) | Whole midbrain | 28195 | 2242 | 4592 |
| Wang et al. (2021) | SN | 39786 | 1259 | 2010.5 |
| Welch et al. (2019) | SN | 64441 | 934 | 1304 |

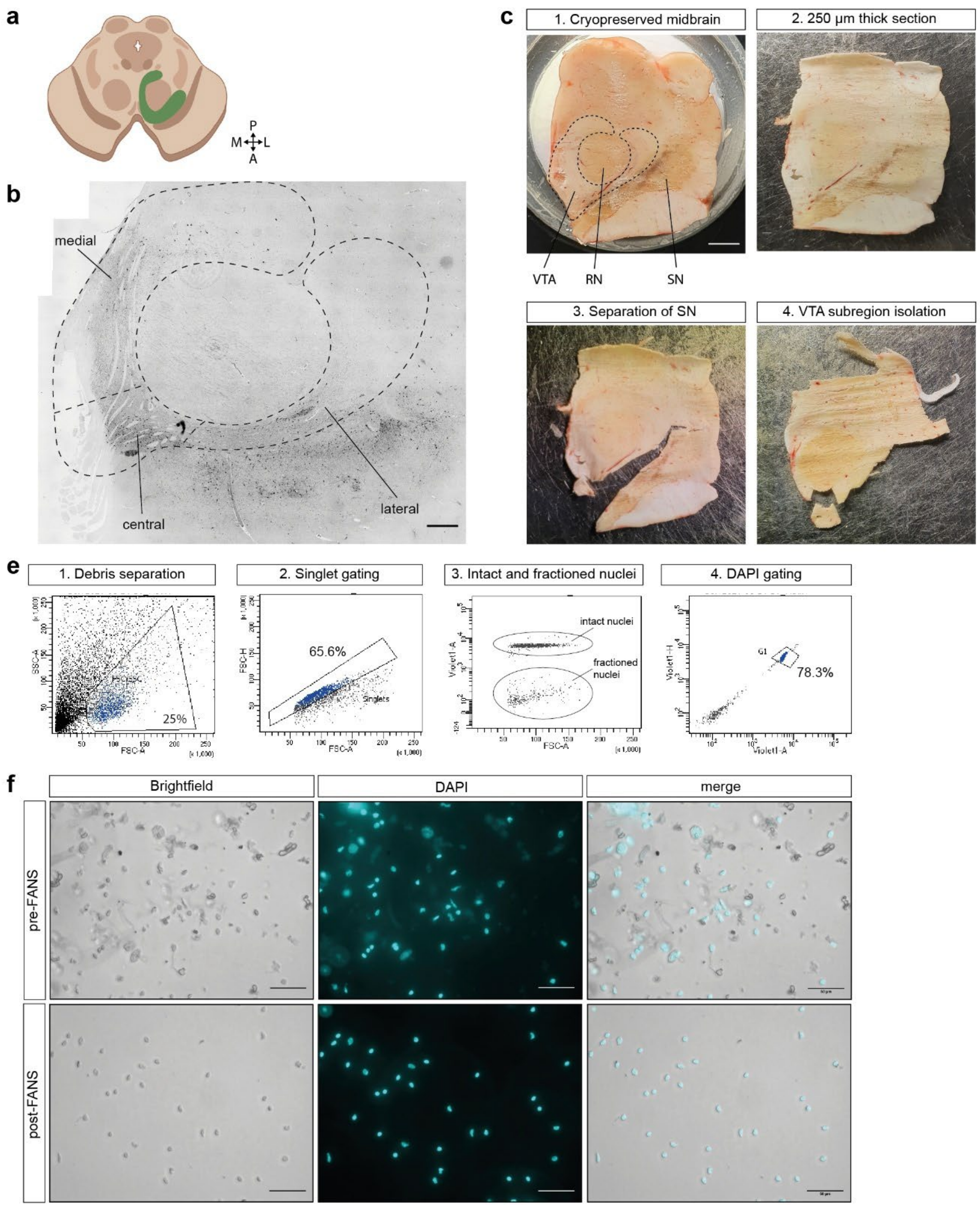

**Supplementary Fig. 1 - Nuclei isolation from the human VTA. a**, Illustration of the ventral tegmental area (VTA) in the human midbrain with anatomical coordinates. A, anterior; P, posterior; M, medial; L, lateral. **b**, Tyrosine hydroxylase (TH) immunohistochemistry staining image of the human midbrain of a non-demented control donor (NBB 2000-036) with three VTA subregions indicated. Scale bar = 1 mm. **c**, Representative images of the VTA tissue isolation. First, the location of the VTA, red nucleus (RN), and substantia nigra (SN) was identified. Then, 250 μm-thick midbrain tissue sections were cut on a cryostat. Per section, the pigmented area of the SN was cut off with a pre-cooled scalpel. Then, three different regions of the VTA were cut out: (i) the lateral part located between the SN and RN, (ii) the medial part located between the midline and the red nucleus and the area more anterior to the red nucleus, and (iii) the part located centrally at the base of the medial and lateral parts where the oculomotor nerve (CN III) enters the midbrain. **e**, Representative images of the fluorescent activated nuclei sort (FANS). **f**, Representative images of DAPI-positive nuclei pre- and post-FANS. Scale bar = 50 μm.

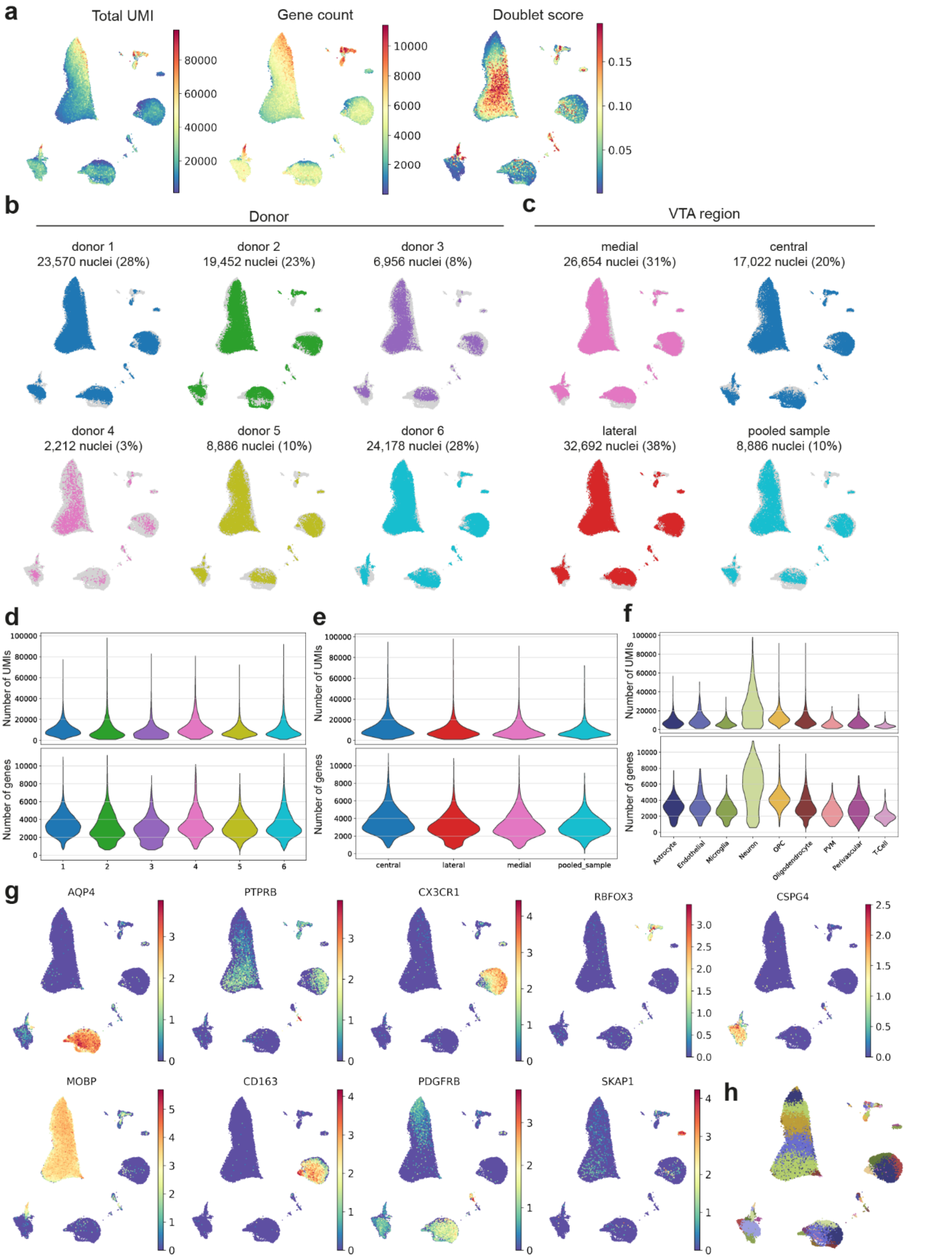

**Supplementary Fig. 2 - Quality control and cell class annotation of the human VTA snRNA-seq dataset. a**, UMAP representations of total UMI, gene count, and doublet score in the snRNA-seq dataset after quality control filtering. **b**, UMAPs of the six donors and the number of nuclei per donor. **c**, UMAPs of the VTA regions and the number of nuclei per region. **d-f**, Violin plots showing the number of UMIs and genes per donor (**d**), VTA region (**e**), and cell class (**f**). **g**, UMAP of normalized cell class-specific marker gene expression. **h**, UMAP of the 68 clusters.

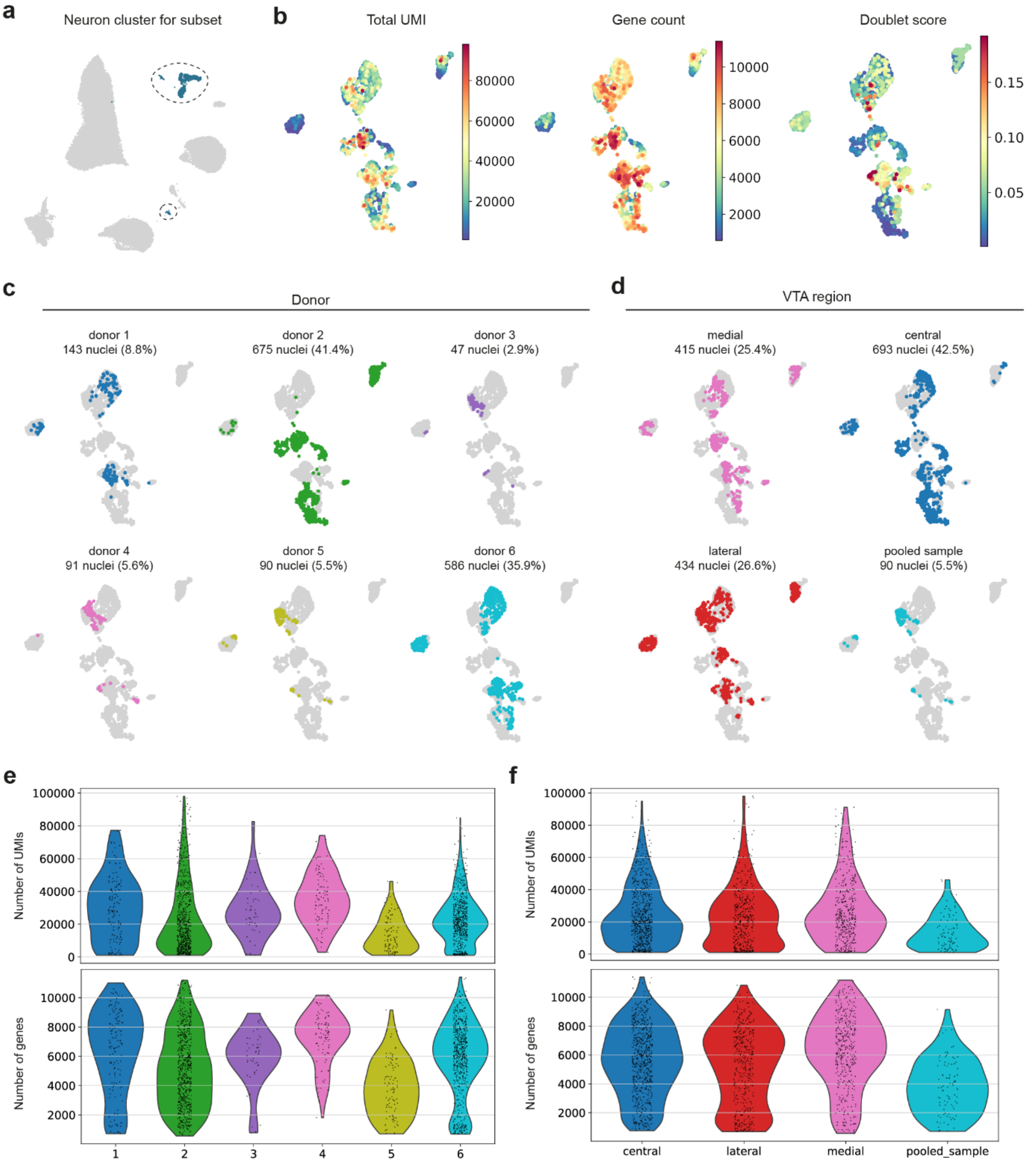

**Supplementary Fig. 3 - Donor and VTA region contributions to the human VTA neuronal cell class. a**, UMAP of the VTA with the selected cluster for further in-depth analysis of neurons. **b**, UMAPs of total UMI, number of genes, and doublet score in the neuronal nuclei. **c-d**, UMAP representations of nuclei contribution per donor (**c**) and VTA region (**d**). **e-f**, Violin plots of the number of UMIs and genes per donor (**e**) and VTA region (**f**).

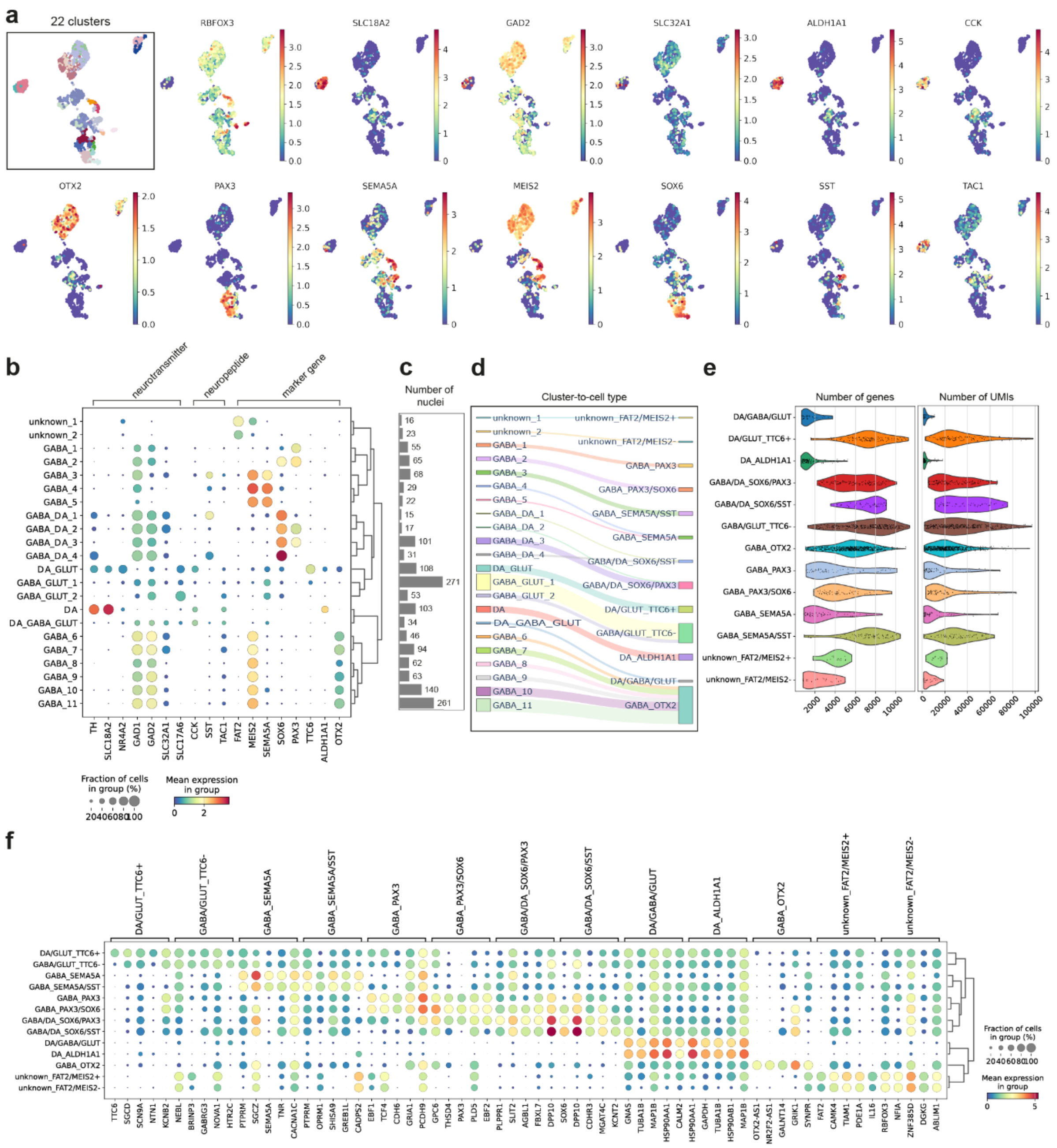

**Supplementary Fig. 4 - Gene expression profiling and quality control of VTA neuronal cell types with snRNA-seq. a**, UMAPs showing the 22 clusters and normalized gene expression for subclass- and cell type-specific genes. **b**, Dot plot of normalized gene expression profiles per neuronal cluster, classifying neurons by pan-neuronal markers, neurotransmitters, neuropeptides, and cell type-specific marker genes. The sizes of the dots indicate the cellular fraction within a cluster that expresses a gene, and colors indicate the mean gene expression levels in this fraction. **c**, The number of nuclei per cluster. **d**, Sankey diagram depicting the mapping of the clusters to the annotated cell types, with line width scaled proportionally to nuclei count.**e**, Violin plots of the number of genes and UMIs per neuronal cell type. **f**, Dot plot of the five most differentially expressed cell type-specific genes.

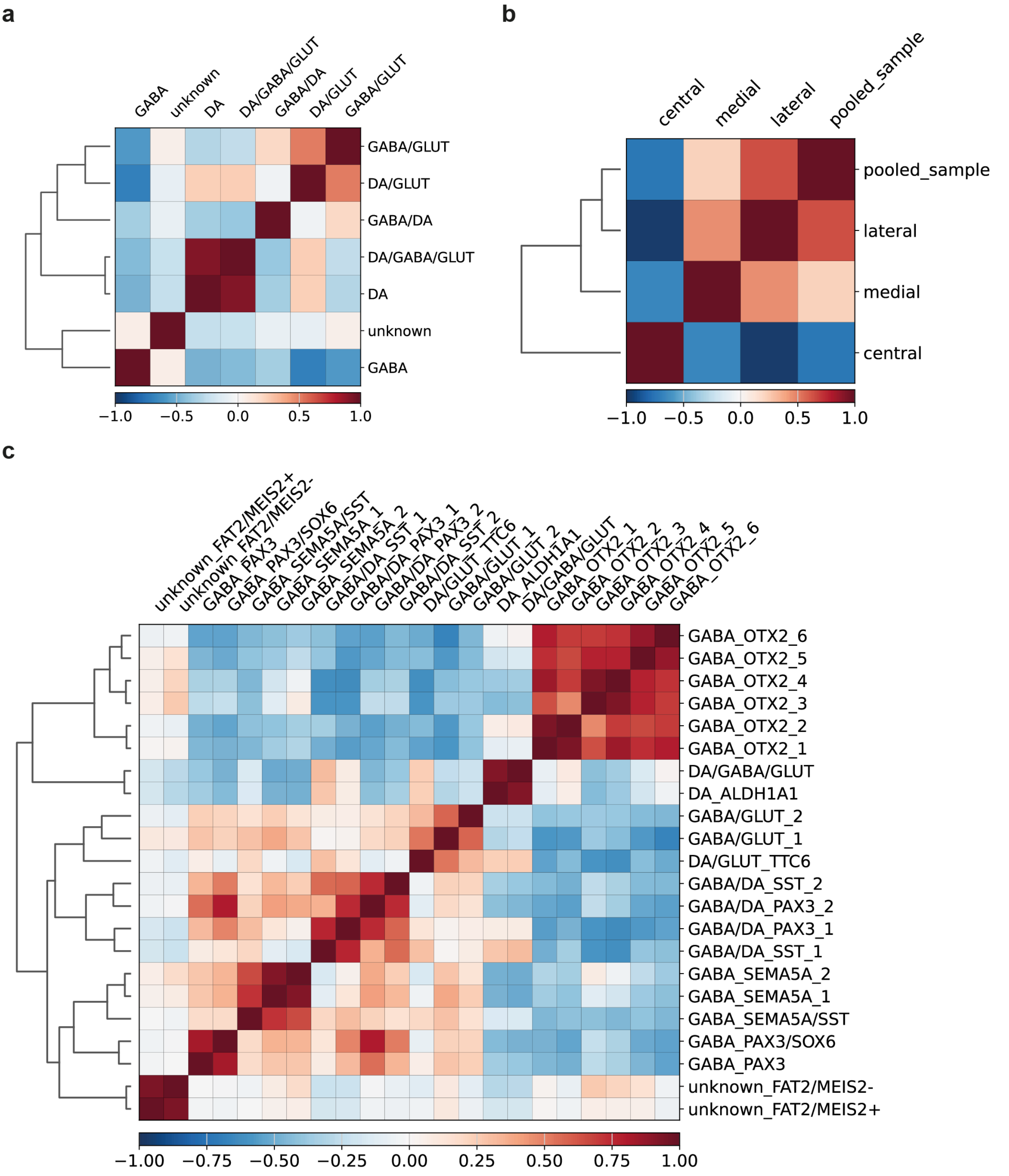

**Supplementary Fig. 5 - Correlation of human VTA neurons. a-c**, Correlation of neurons between subclasses (**a**), VTA regions (**b**), and cell type-annotated clusters (**c**).

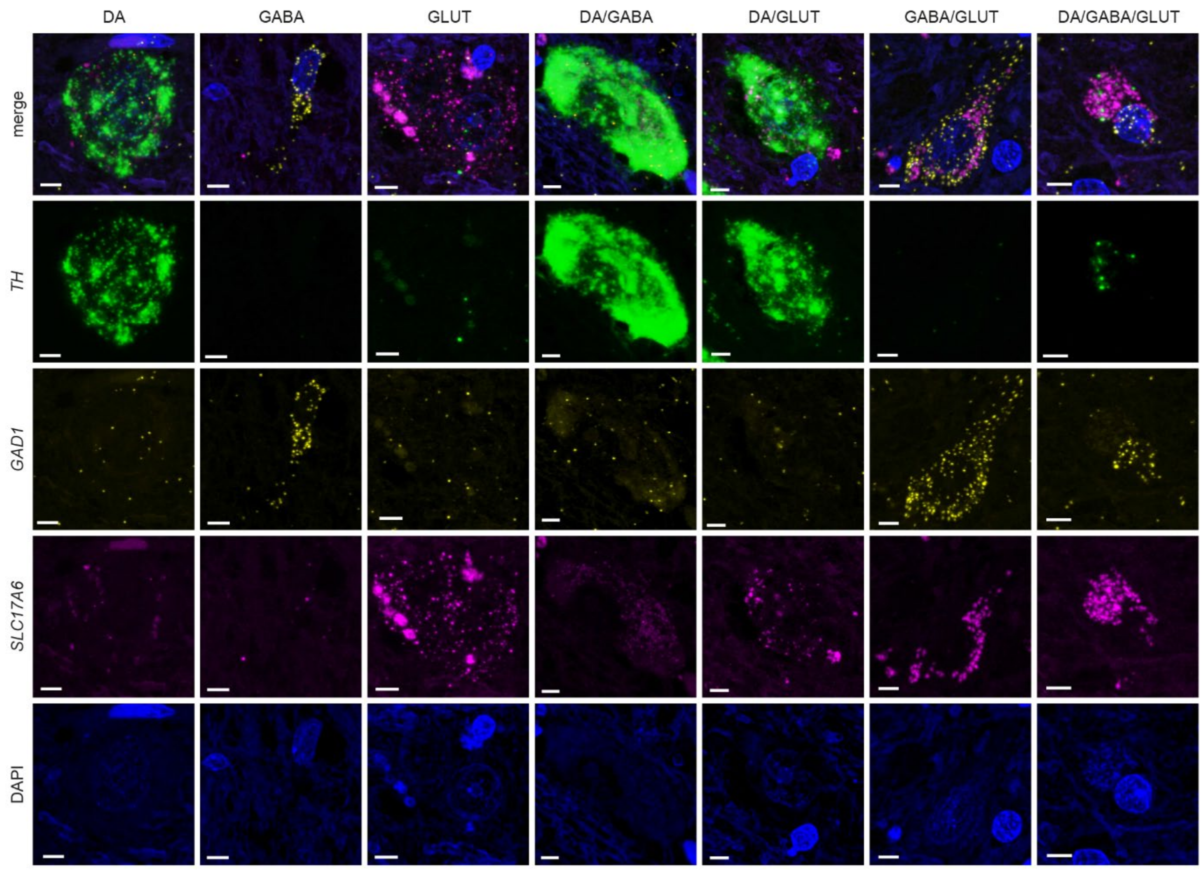

**Supplementary Fig. 6 - Validation of the seven neuronal subclasses *in situ* with RNAscope.** Representative images from the detected signal from neurotransmitter-specific genes *TH* (dopaminergic), *GAD1* (GABAergic), and *SLC17A6* (glutamatergic) in the VTA of a non-demented control donor. The images are maximum intensity projections of Z-stacks. Donor 2000-036. Scale bar = 5 μm.

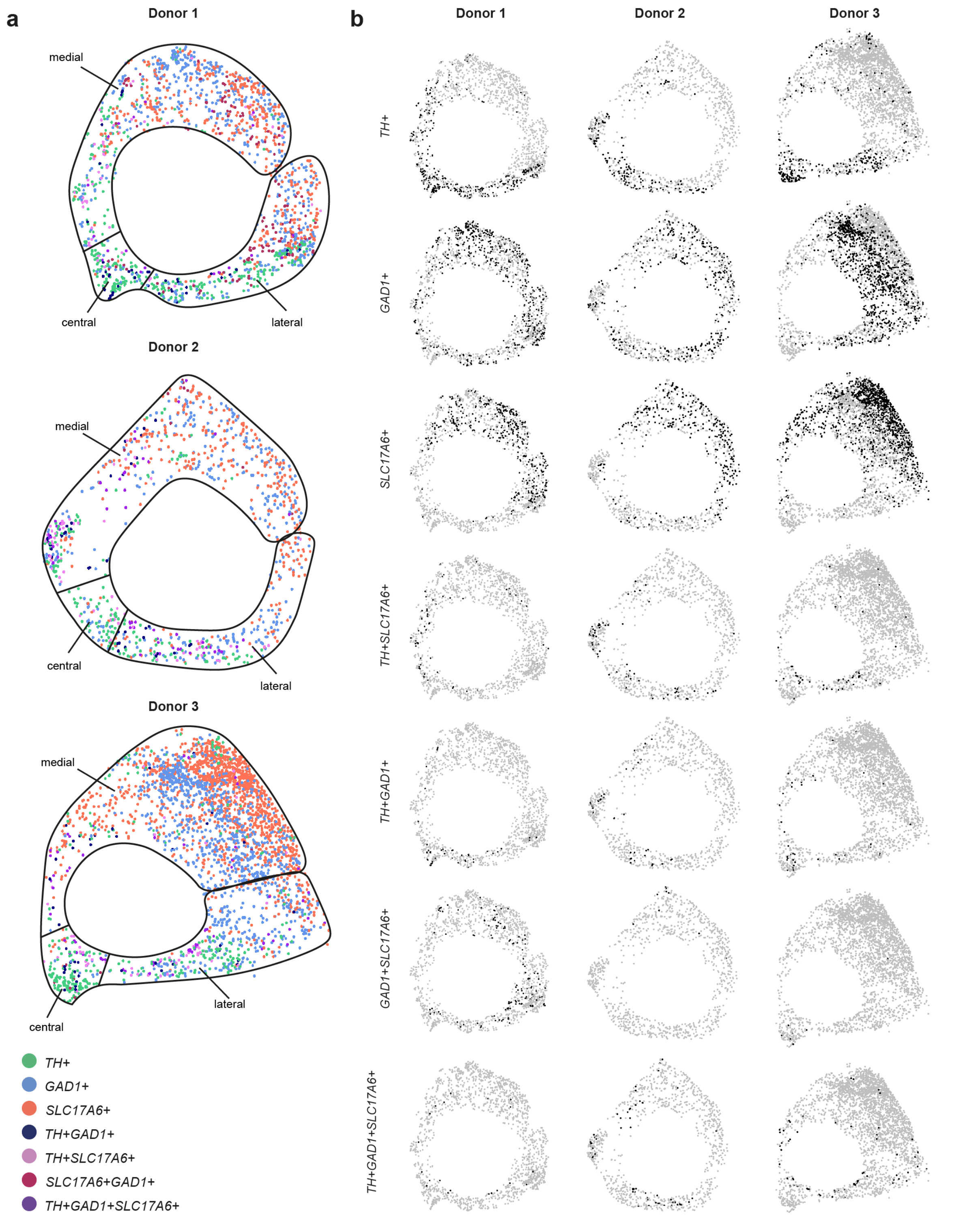

**Supplementary Fig. 7 - Spatial localization of the neuronal subclasses in the human VTA with RNAscope. a**, Prevalence of all neuronal subclasses in a tissue section of three non-demented control donors. Donor 1: 2000-036, donor 2: 1998-104, donor 3: 1999-101. **b**, Localization of individual subclasses in each donor.

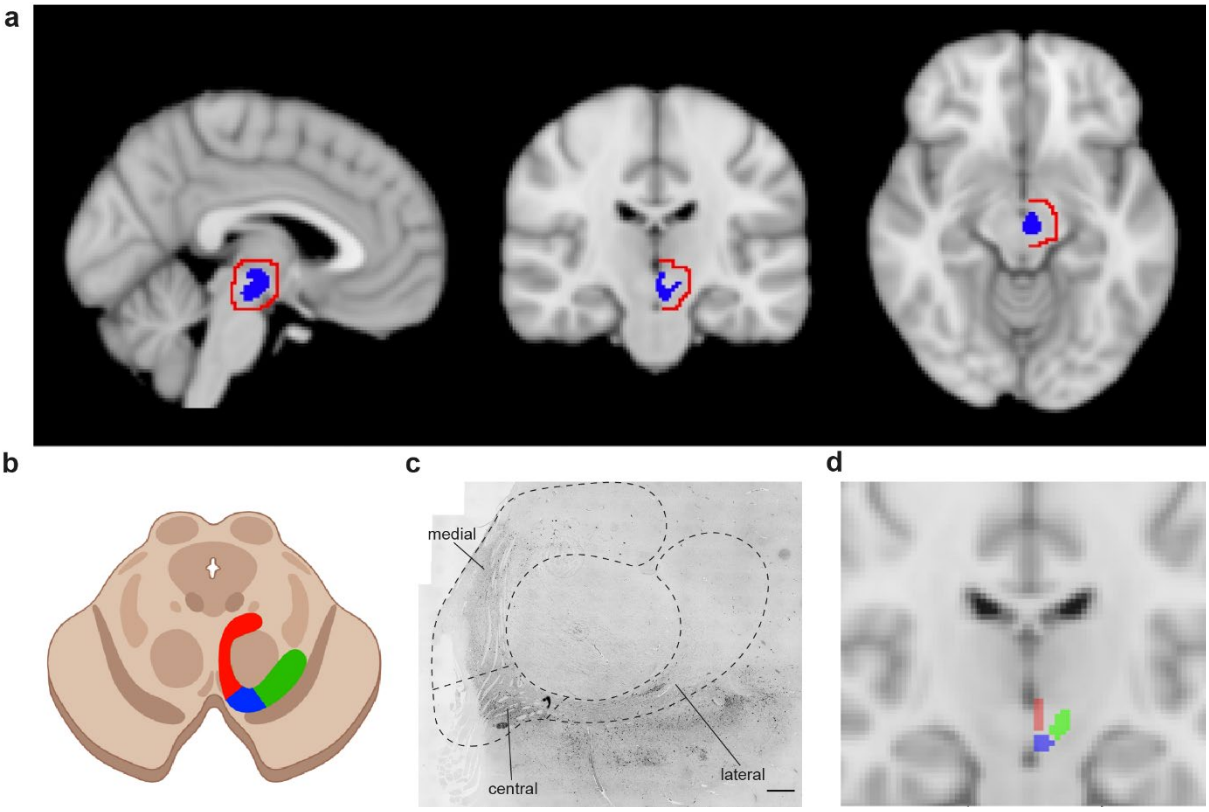

**Supplementary Fig. 8 – VTA segmentation for regional connectivity analysis with fMRI. a,** In blue is the original atlas of the left VTA with the threshold set at 0.1. In red is the mask of the vicinity of the VTA, for which the average signal was included as a covariate in the seed-based analysis. Note that the mask was restricted to the left hemisphere. **b,** Schematic of the human midbrain with color-coded VTA subregions. **c**, Microscopy image of the VTA. Dashed lines represent the boundaries of the dissected subregions for snRNA-seq. Scale bar = 1 mm. **d**, The manual segmentation of the VTA regions for fMRI analysis. Medial, central, and lateral regions are shaded in red, blue, and green, respectively.

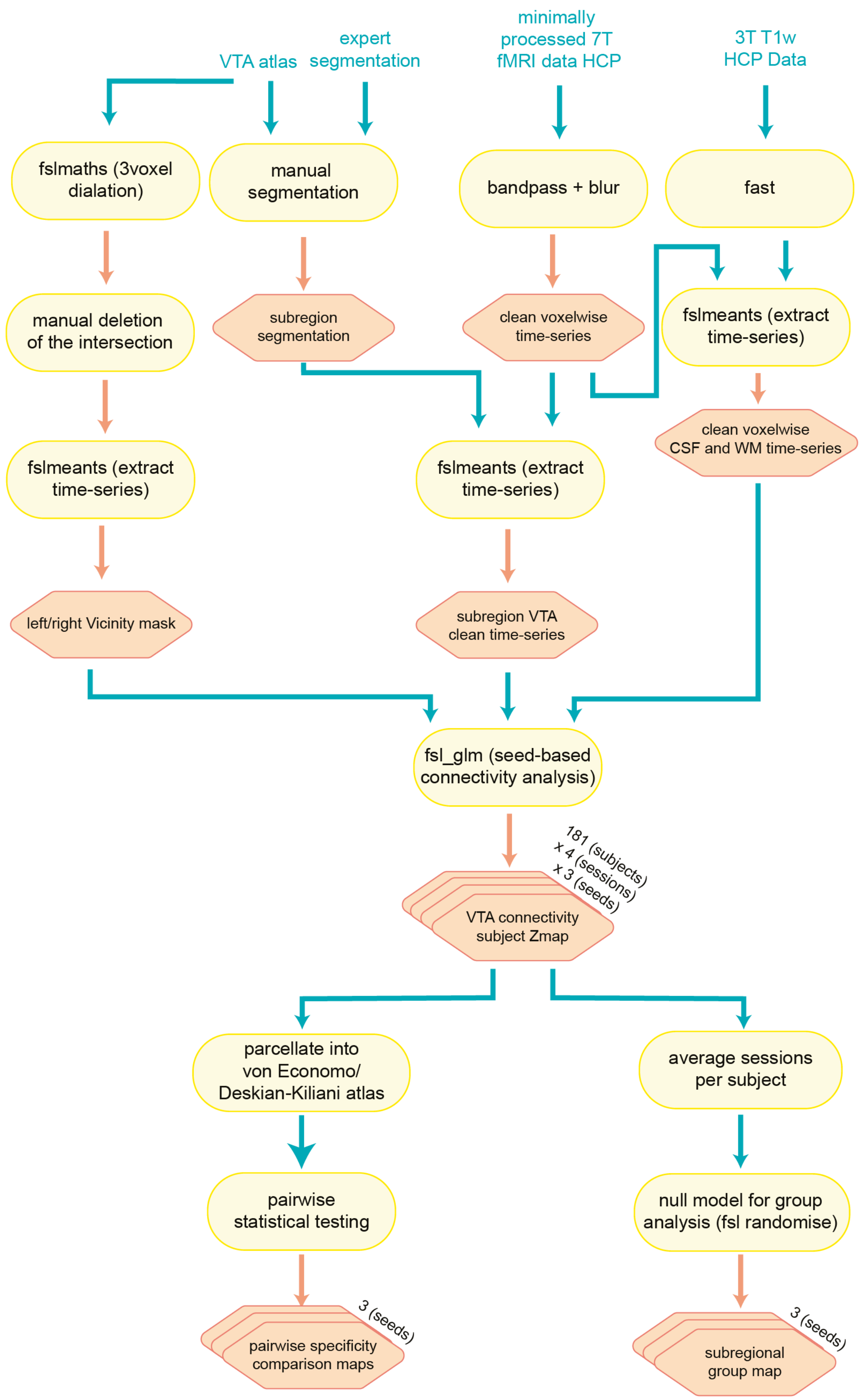

**Supplementary Fig. 9 - fMRI analysis workflow.**

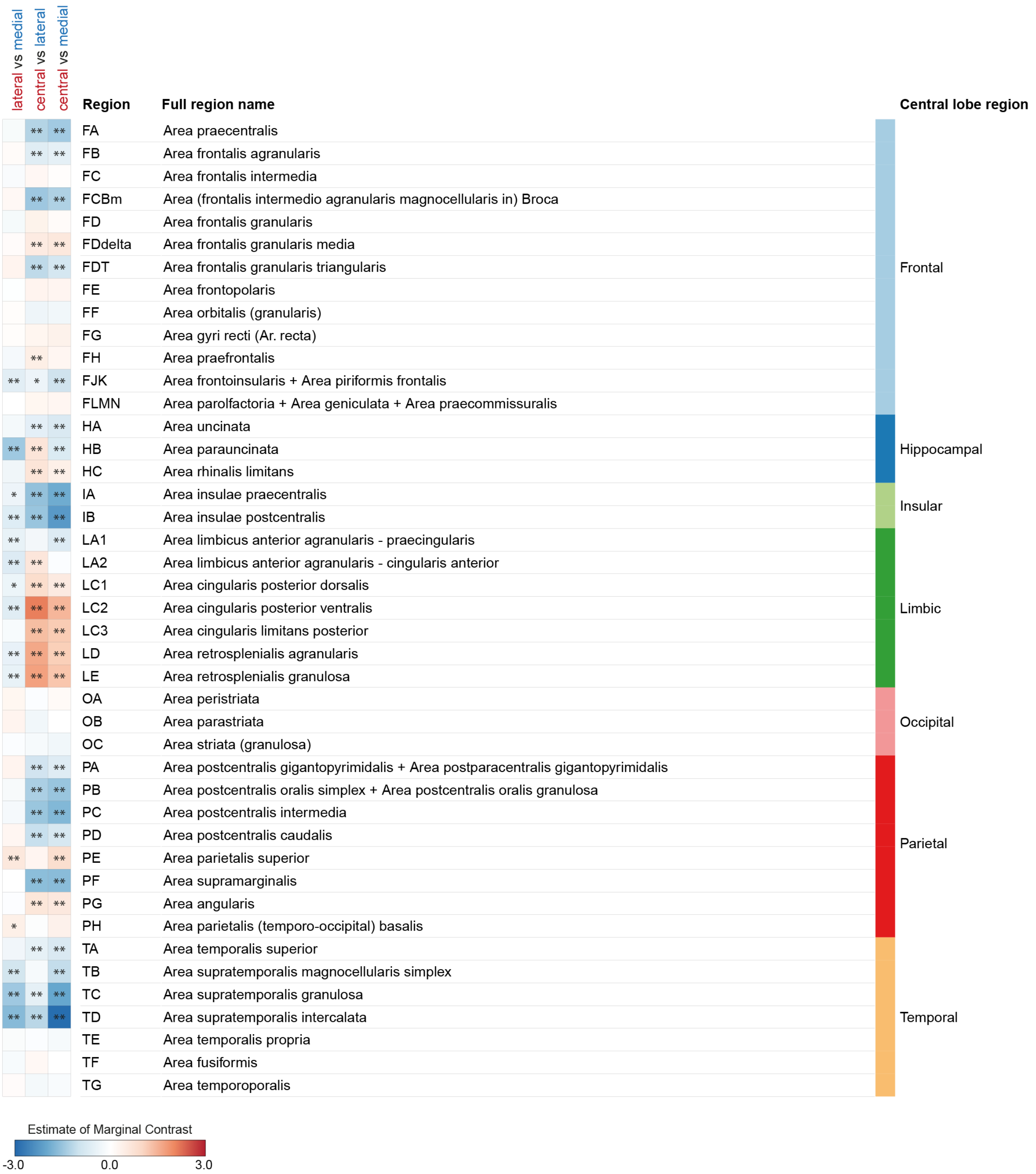

**Supplementary Fig. 10 – VTA subregions show different connectivity profiles using the Von Economo–Koskinas MRI-based cortical parcellation.** Pairwise analyses for region-specific connectivity. Each column shows a different pair of VTA subregion comparisons for every parcellated brain region (rows). The connectivity strength for each comparison is color-coded and shown in t-values, and its significance is indicated by *= p<0.1, **=p<0.05, and ***= p<0.01. Analysis details are further described in **Suppl. Note N1**. Region indicates the Von Economo–Koskinas abbreviations.

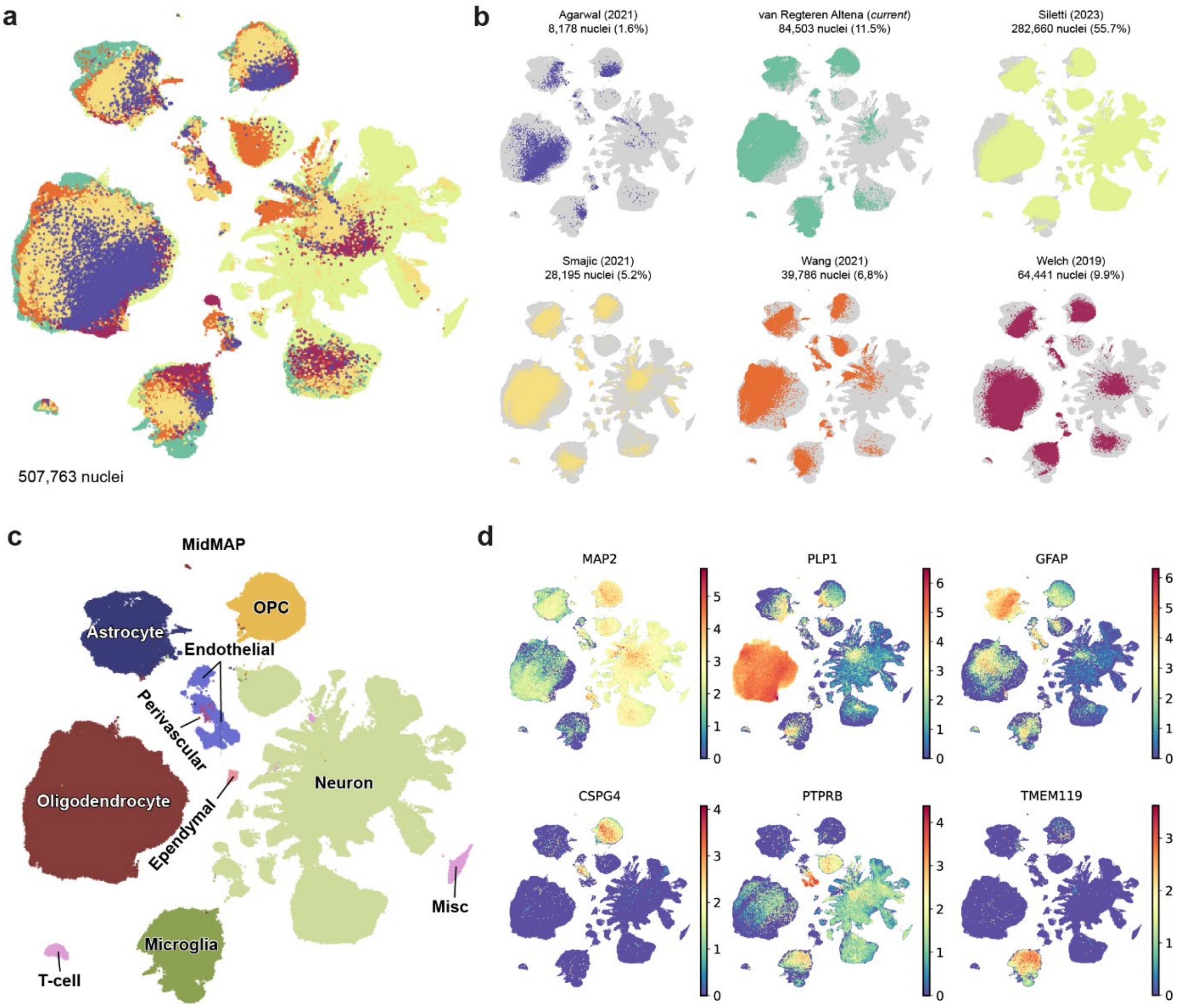

**Supplementary Fig. 11 - Integration of human midbrain snRNA-seq datasets. a,** UMAP of 507,763 integrated nuclei from six independent snRNA-seq adult human midbrain datasets. **b,** UMAPs of the six datasets and the number of nuclei contributing to the integrated midbrain dataset. **c,** UMAP of the integrated dataset annotated with the k-nearest neighbors predicted cell type. **d**, UMAPs showing normalized gene expression of cell class-specific gene expression.

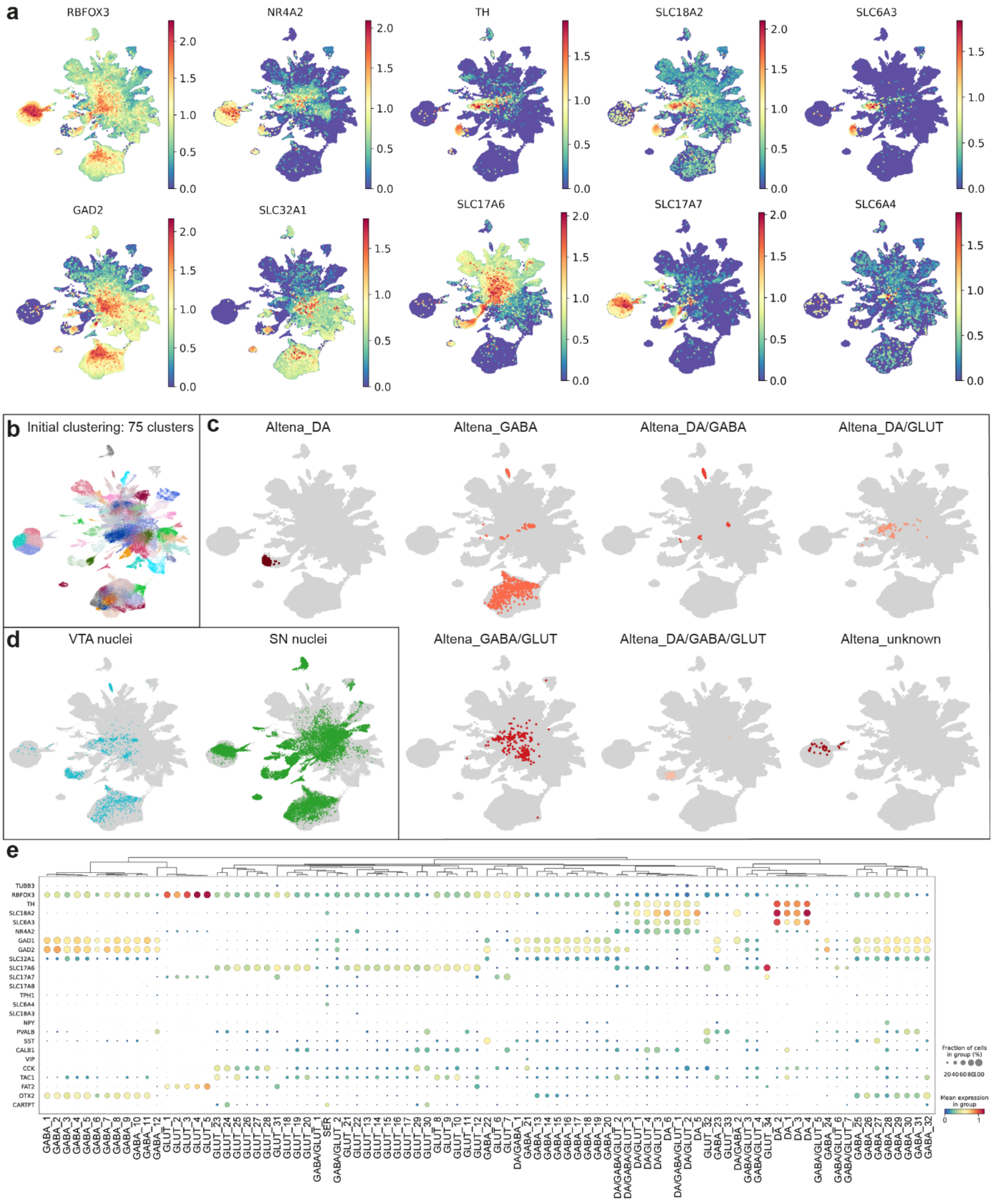

**Supplementary Fig. 12 - Neuronal subcluster analysis of the integrated human midbrain atlas. a**, Normalized gene expression profiles of neuronal and neurotransmitter-specific markers. **b**, UMAP of the initial 75 identified neuronal clusters. **c**, The neuronal subclass annotated nuclei from the human VTA atlas, plotted on the UMAP of the human midbrain neuronal atlas. **d**, UMAP plotting of VTA- and SN-identified nuclei within the total neuronal cell class population of the human midbrain. **e**, Dot plot of neuronal marker genes to identify the hierarchically-organized neuronal clusters.

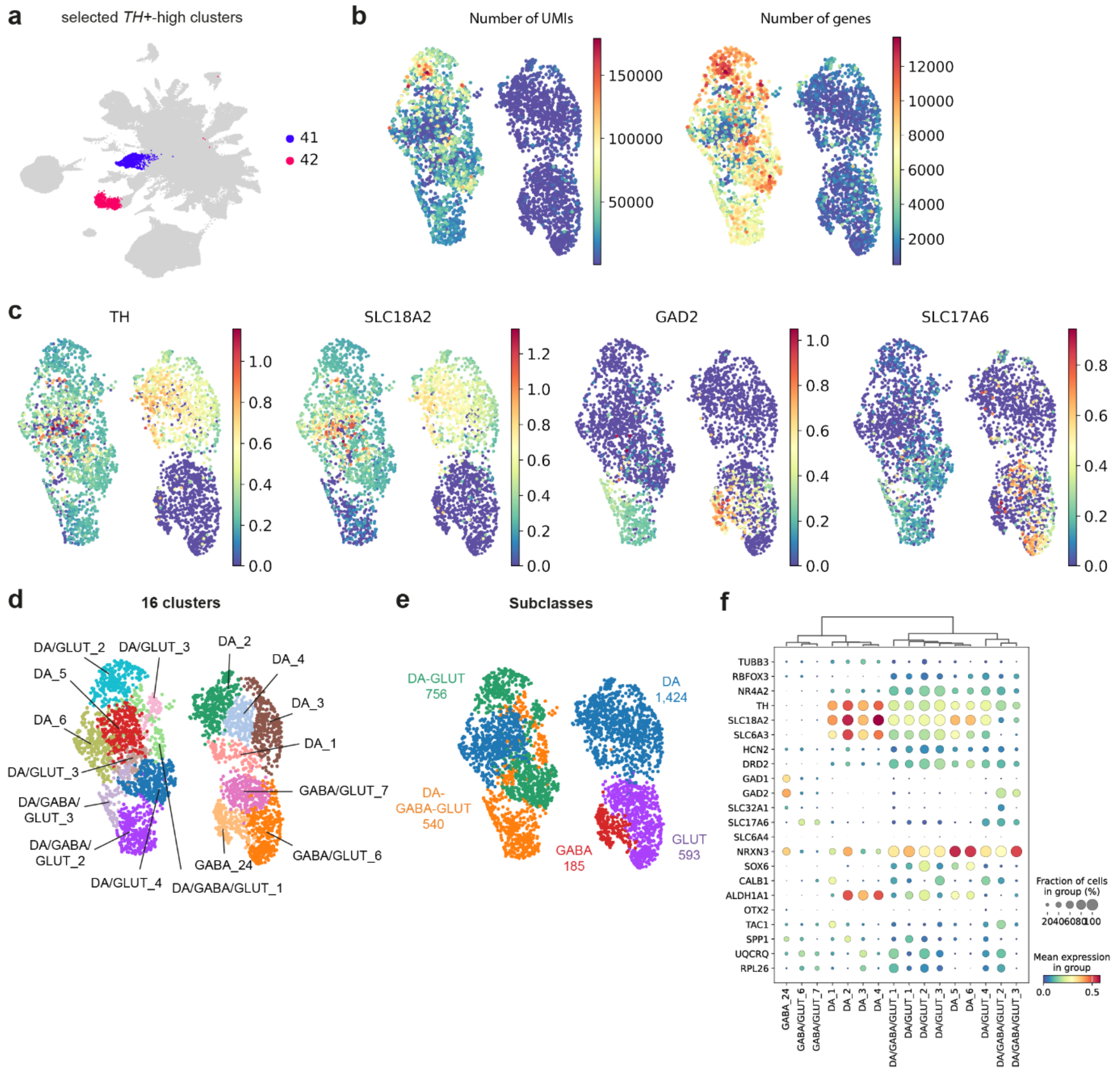

**Supplementary Fig. 13 - Identification of DA neuronal subclass clusters using the integrated midbrain atlas. a**, The two *TH*-high clusters (41 and 42) that were isolated to identify the DA neuron cell type profiles in the human midbrain in more detail. **b**, Quality control metrics number of UMIs and number of genes in clusters 41 and 42. **c**, Normalized expression of neurotransmitter-specific marker genes. **d**, UMAP of the 16 identified clusters with Leiden (resolution 1) clustering. **e**, Annotated neuronal subclasses. **f**, Dot plot of marker gene expression per cluster to identify neuronal subclasses.

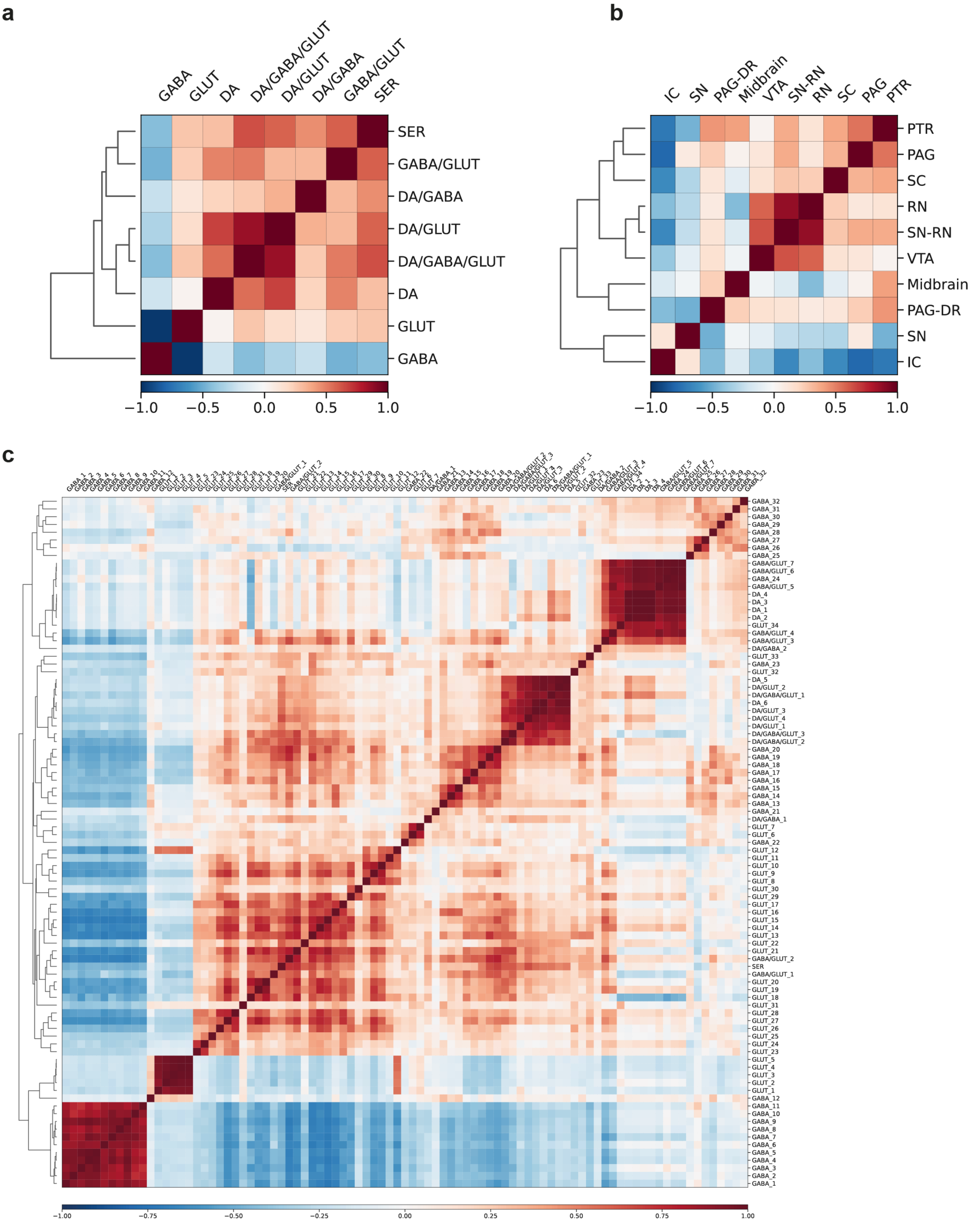

**Supplementary Fig. 14 - Correlation of midbrain neurons. a-c**, Correlation of neuronal subclasses (**a**), midbrain regions (**b**), and clusters (**c**) within the midbrain.

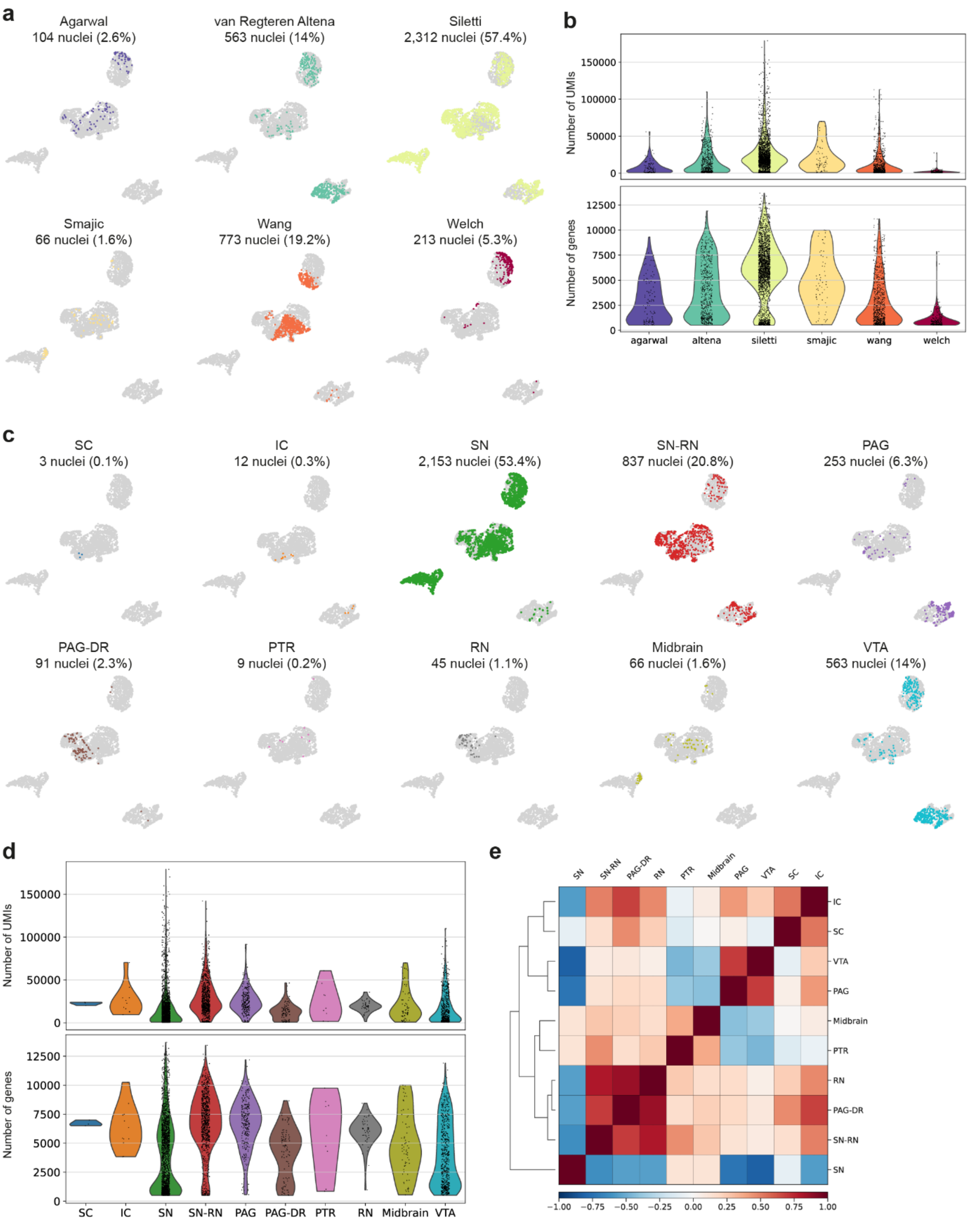

**Supplementary Fig. 15 - Atlas and midbrain region distributions of the DA neuronal clusters. a**, UMAPs of the individual atlases. **b**, Violin plots of the number of UMIs and the number of genes per atlas. **c**, UMAPs of the different midbrain regions. **d**, Violin plots of the number of UMIs and the number of genes in the DA subclasses per nucleus per midbrain region. **e**, Correlation matrix of the different midbrain regions.

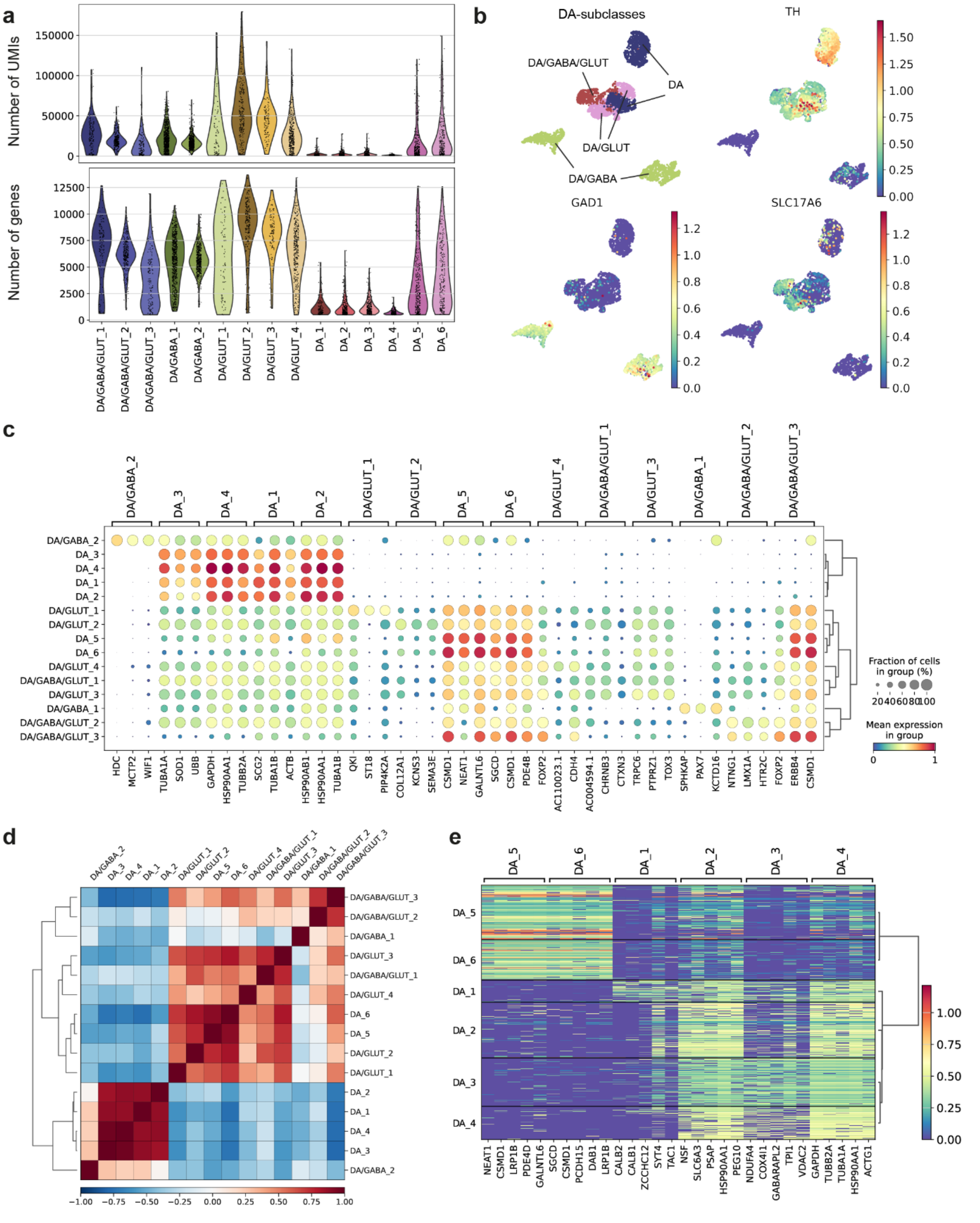

**Supplementary Fig. 16 - Identification of DA cluster gene profiles. a**, Violin plots of the number of UMIs and the number of genes detected in the DA clusters. **b**, UMAPs showing the DA subclass annotations and normalized gene expression profiles of genes used for their identification. **c**, Dot plot of the top three differentially expressed genes per DA cluster. **d**, Correlation matrix of the clusters. **e**, Heatmap of the top five differentially expressed genes of each DA-only cluster, shown as normalized expression per nucleus in each.

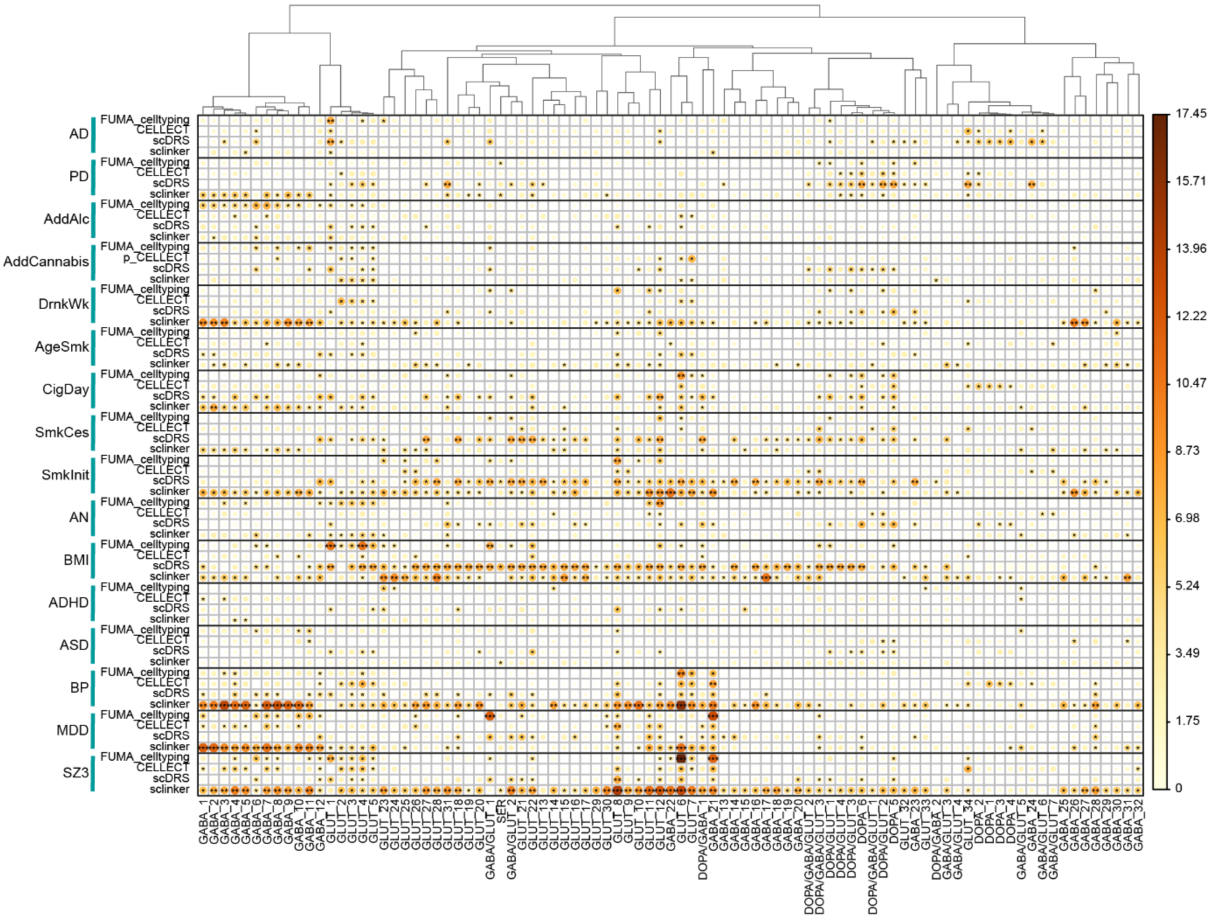

**Supplementary Fig. 17 – Neuronal cell type prioritization in the human midbrain per method for each tested trait.** The color of the dots indicates the p-value for the association per cell type. Asterisks indicate the significance of the minimal p-value for each method, after multiple testing correction for the number of cell types: * = marginal significance (0.05), ** = significance within the phenotype, corrected for all cell types (0.05/89), *** = significance across phenotypes and cell types (0.05/89*16). AD, Alzheimer's disease; PD, Parkinson's disease; AddAlc, alcohol addiction; AddCannabis, cannabis addiction; DrnkWk, drinks per week; AgeSmk, age smoking; CigDay, cigarettes per day; SmkCes, ever stopped smoking; SmkInit, ever smoked; AN, anorexia nervosa; BMI, body mass index; ADHD, attention deficit/hyperactivity disorder; ASD, autism spectrum disorder; BP, bipolar disorder; MDD, major depressive disorder; SZ3, schizophrenia.

### Supplementary Note 1 – Functional Magnetic Resonance Analyses

#### Additional Methods

*Preprocessing of (f)MRI*

Structural MRI data were acquired on a customised Siemens 3T "Connectome Skyra", using a standard 32-channel Siemens receive head coil, resulting in images with 0.7 mm isotropic voxels. Functional MRI data were acquired on a Siemens Magnetom 7T MR scanner, using a Nova32 32-channel Siemens receive head coil with an incorporated head-only transmit coil surrounding the receive head coil from Nova Medical. The data were already minimally preprocessed by the Human Connectome Project team. Reference images were aligned to the structural scan and resampled while preserving the resolution from the native space of the fMRI acquisitions. Next, intensity normalisation and bias field removal were performed. Finally, temporally filtering using a 2000s high-pass filter was applied, after which the data was automatically denoised using FIX. We applied an additional high-pass filter (cut-off frequency = 0.1Hz) to the data.

*Covariate Residualisation*

We segmented whole-brain, white matter, grey matter, and cerebrospinal fluid (CSF) on the skull-stripped T1-weighted image using FSL FAST[1,2]. The time series of voxels in the whole-brain, white matter, and CSF masks were averaged for each session of each subject using fslmeants. Additionally, the average time series of a 1-voxel-thick mask (at a distance of three voxels) around the left VTA was also regressed to improve the signal by controlling further for physiological noise in the vicinity of the VTA (**Suppl. Fig. 6a**). The covariate residualisation was performed per subject session using FSL FEAT, resulting in 736 clean images (181 subjects * 4 sessions).

*Subnetwork definition*

The definition of resting-state networks (RSNs) based on the Yeo-Krienen 7-network atlas was carried out as described before. Briefly, the Yeo-Krienen atlas contains a parcellation map of 7 large-scale functional resting-state networks, including the visual, somatomotor, dorsal attention, ventral attention (VAN; also referred to as salience), limbic, frontoparietal (FPN; also referred to as central executive), and default mode network (DMN). An annotation file of the 7 functional networks was included for the fsaverage subject in the FreeSurfer Software package (https://surfer.nmr.mgh.harvard.edu/fswiki/CorticalParcellation_Yeo2011). For this, the surface-based annotation was translated to a 3D brain volume in volumetric space, in which each grey matter voxel was assigned a network label. Next, for each region in the Desikan-Killiany atlas, the ratio of voxels that belonged to each of the 7 RSNs was computed. Using a majority vote approach, the label of the functional network corresponding to most voxels was then assigned to that region. All the regions in the aseg atlas were assigned to the subcortical network.

#### Replication with Von Economo-Koskinas Atlas

*Registration*

We registered the Von Economo-Koskinas cortical atlas with the 1.6mm resolution MNI152 template space using FSL MCFLIRT (nearest neighbor interpolation, 12 degrees-of-freedom)[3]. This atlas comprises 84 cortical areas, 42 on each cerebral hemisphere. The statistical analysis was restricted to the left hemisphere to align with the experimental data. Our atlas was applied to the functional connectivity maps of each subject for the 4 sessions to calculate the average functional connectivity between each of the three VTA regions and each of the 42 left cortical areas. Cortical areas for which the connectivity value deviates more than 3 standard deviations from the mean connectivity of that region in all subjects per session were considered outliers and removed from the analysis.

*Statistical Analyses*

For subsequent statistical analyses using the Von Economo-Koskinas atlas, the threshold for significance was adjusted to correct for multiple comparisons, factoring in the number of seeds (3), cortical regions (42) in the left hemisphere, and interactions (42*3) (alpha = 0.05/(3 + 42 + 3*42) = $2.9 \cdot 10^{-4}$). Statistical analysis on this atlas shows a large overlap with the Desikan-Killiany Atlas. The results are summarized in **Suppl. Fig. 8.** For example, the area supertemporalis granulosa (TC) and area supertemporalis intercalata (TD) in the Von Economo-Koskinas atlas show anatomical overlap with the transverse temporal region in the Desikan-Killiany Atlas, which both show the strongest connectivity with the medial VTA.

**Supplementary Note 2 - GWAS to cell type analyses**

*GWAS Preprocessing*

We applied GWAS to cell type methods for 16 phenotypes related to neuropsychiatric disorders (Methods). In the first step, we preprocessed all summary statistics using MAGMA[4] (version 1.10) to aggregate SNP-level genetic signal into gene-based Z-scores using the multi=snp-wise flag. SNPs were assigned to a gene if they were between the transcription start and stopping sites. In addition, the summary statistics were preprocessed with the munge_sumstats.py script that is part of the linkage disequilibrium score regression (LDSC) pipeline[5].

*GWAS to cell type methods*

Methods that prioritize cell types associated with complex traits based on GWAS differ in their assumptions of what genes are used to identify a cell type. For this reason, we applied several methods to identify midbrain neuronal cell types associated with neuropsychiatric traits. See below for details on assumptions and implementation.

*FUMA cell typing*

FUMA cell typing assumes that (relative) mean expression of genes should correlate with GWAS signal[6]. To implement this strategy, mean expression per cell type, and average expression over all cell types were calculated per gene. Mean expression was then associated with the MAGMA-derived Z-scores using gene property analysis implemented in MAGMA[4] using overall average expression as a covariate.

*CELLECT*

CELLECT is a toolkit that is based on the specificity of genes in cell types[7]. Specificity of a gene can be interpreted as a measure of how much a gene is expressed in a certain cell type compared to other cell types. CELLECT computes four different specificity indices, after which the average of those is used as a continuous annotation in LD score regression. It includes the standard baseline annotations for LDSC regression version 1.1, the cell type annotation, and an additional annotation including all SNPs in genes in the VTA dataset. This framework uses HapMap 3 SNPs excluding the MHC region, and the 1000 Genome version 3 as LD reference panel.

*scDRS*

The first step in the scDRS pipeline[8] is to identify the genes that carry the largest signal in the GWAS, which is done by selecting the top 1000 genes based on MAGMA Z-scores. For these genes, an aggregated expression score is computed for each cell, inversely weighted by an estimate of technical noise. Statistical significance is estimated using a permutation procedure, comparing the aggregated expression score to a number of so-called control genes: randomly selected genes matched for mean expression and expression variance, but not associated in the GWAS. The top 5% quantile of the aggregated scores per cell type is used as the test statistic for cell type specificity. scDRS was implemented following instructions on https://martinjzhang.github.io/scDRS/. The default setting for the number of scDRS does not allow for p-values smaller than our multiple testing threshold within a phenotype (see below); we adjusted the number of permutations to 25000.

*sc-linker*

sc-linker selects genes representative of cell types by first running a nonparametric Wilcoxon's rank-sum test to test differential expression for each gene[9]. A SNP-based annotation per gene is created using an enhancer-gene link strategy based on known Activity-by-contact and tissue-specific based on the

Roadmap Epigenetics Consortium data-specific annotation was added to the baseline annotation categories of LDSC, version 2.1[10,11]. The original sc-linker pipeline uses an empirical distribution to create p-values, but since we have a limited number of cell types this did not seem appropriate here. Instead, p-values for cell types were taken as the p-values for the tau-coefficient as suggested in Finucane et al., 2015[5]. The sc-linker implementation was based on the github repositories https://github.com/karthikj89/scgenetics and https://github.com/kkdey/GSSG.
